# Manipulation of the unfolded protein response: a pharmacological strategy against coronavirus infection

**DOI:** 10.1101/292979

**Authors:** Liliana Echavarría-Consuegra, Georgia M. Cook, Idoia Busnadiego, Charlotte Lefèvre, Sarah Keep, Katherine Brown, Nicole Doyle, Giulia Dowgier, Krzysztof Franaszek, Nathan A. Moore, Stuart G. Siddell, Erica Bickerton, Benjamin G. Hale, Andrew E. Firth, Ian Brierley, Nerea Irigoyen

## Abstract

Coronavirus infection induces the unfolded protein response (UPR), a cellular signalling pathway composed of three branches, triggered by unfolded proteins in the endoplasmic reticulum (ER) due to high ER load. We have used RNA sequencing and ribosome profiling to investigate holistically the transcriptional and translational response to cellular infection by murine hepatitis virus (MHV), often used as a model for the *Betacoronavirus* genus to which the recently emerged SARS-CoV-2 also belongs. We found the UPR to be amongst the most significantly up-regulated pathways in response to MHV infection. To confirm and extend these observations, we show experimentally the induction of all three branches of the UPR in both MHV- and SARS-CoV-2-infected cells. Over-expression of the SARS-CoV-2 ORF8 or S proteins alone is itself sufficient to induce the UPR. Remarkably, pharmacological inhibition of the UPR greatly reduced the replication of both MHV and SARS-CoV-2, revealing the importance of this pathway for successful coronavirus replication. This was particularly striking when both IRE1α and ATF6 branches of the UPR were inhibited, reducing SARS-CoV-2 virion release ∼1,000-fold. Together, these data highlight the UPR as a promising antiviral target to combat coronavirus infection.

**Author Summary:** SARS-CoV-2 is the novel coronavirus responsible for the COVID-19 pandemic which has resulted in over 100 million cases since the end of 2019. Most people infected with the virus will experience mild to moderate respiratory illness and recover without any special treatment. However, older people, and those with underlying medical problems like chronic respiratory disease are more likely to develop a serious illness. So far, more than 2 million people have died of COVID-19. Unfortunately, there is no specific medication for this viral disease.

In order to produce viral proteins and to replicate their genetic information, all coronaviruses use a cellular structure known as the endoplasmic reticulum or ER. However, the massive production and modification of viral proteins stresses the ER and this activates a compensatory cellular response that tries to reduce ER protein levels. This is termed the unfolded protein response or UPR. We believe that coronaviruses take advantage of the activation of the UPR to enhance their replication.

The UPR is also activated in some types of cancer and neurodegenerative disorders and UPR inhibitor drugs have been developed to tackle these diseases. In this work, we have tested some of these compounds in human lung cells infected with SARS-CoV-2 and found that virus production was reduced 1000-fold in human lung cells.

## Introduction

The *Coronaviridae* are a family of enveloped viruses with positive-sense, non-segmented, single-stranded RNA genomes. Coronaviruses (CoVs) cause a broad range of diseases in animals and humans. SARS-CoV, MERS-CoV and SARS-CoV-2, members of the genus *Betacoronavirus*, are three CoVs of particular medical importance due to high mortality rates and pandemic capacity [1–3]. SARS-CoV-2 is the causative agent of the current COVID-19 pandemic, which has resulted in over 95 million cases and more than 2 million deaths since the end of 2019. Although 15% of the cases develop a severe pathology, no specific therapeutic treatment for COVID-19 has been approved to date, highlighting the urgent need to identify new antiviral strategies to combat SARS-CoV-2, besides future CoV zoonoses.

During CoV replication, the massive production and modification of viral proteins, as well as virion budding-related endoplasmic reticulum (ER) membrane depletion, can lead to overloading of the folding capacity of the ER and consequently, ER stress [4]. This activates the unfolded protein response (UPR) which is controlled by three ER-resident transmembrane sensors: inositol-requiring enzyme-1 α (IRE1α), activating transcription factor-6 (ATF6), and PKR-like ER kinase (PERK), each triggering a different branch of the UPR (Fig 1A). Activation of these pathways leads to decreased protein synthesis and increased ER folding capacity, returning the cell to homeostasis [5].

**Figure 1:**
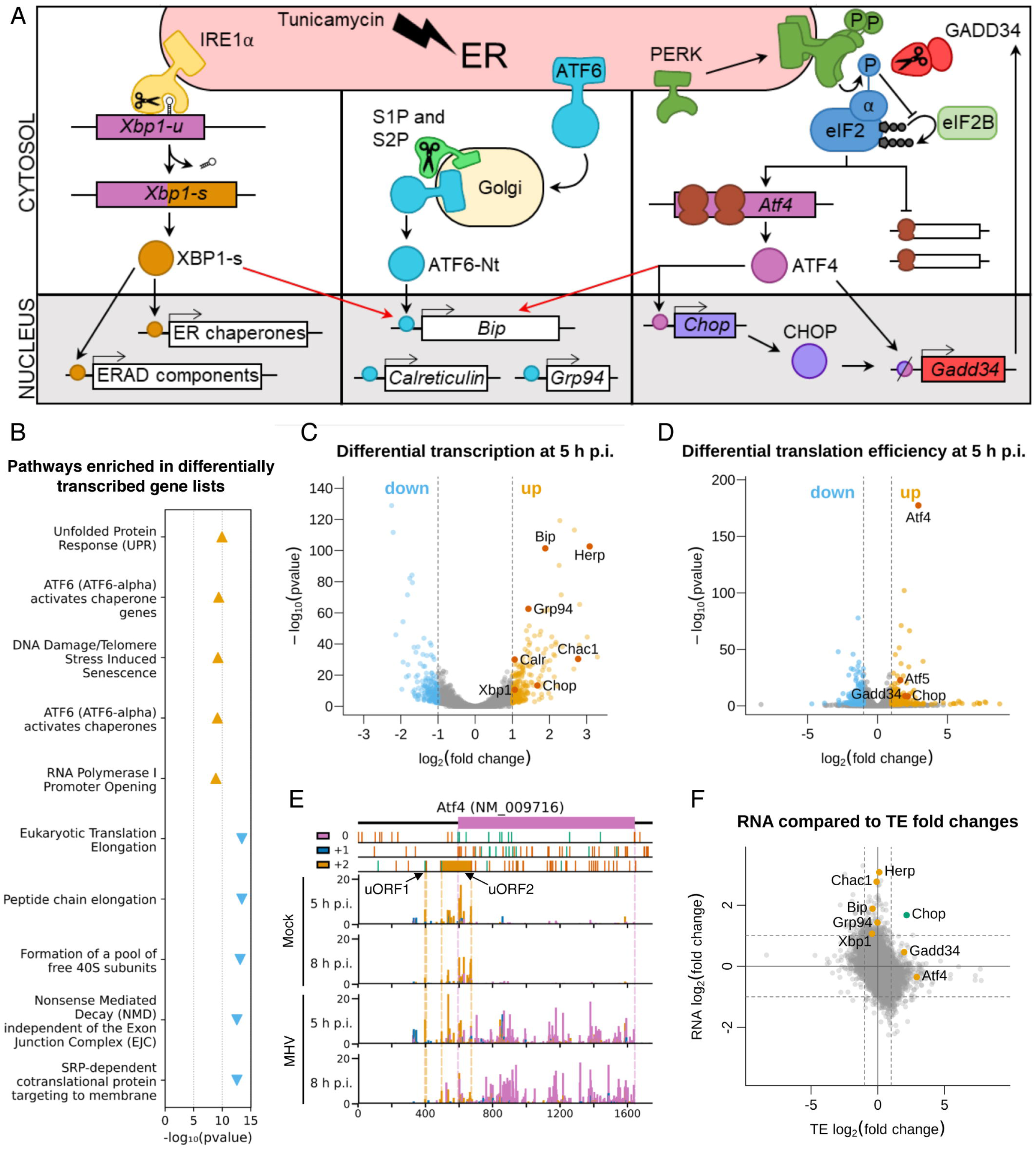
Ribosome profiling reveals the unfolded protein response as a key pathway in the host response to MHV-A59 infection. **(A)** Schematic of the three branches of the UPR (IRE1α, ATF6, and PERK). ERAD = ER-associated protein degradation. **(B)** Top five most significantly enriched Reactome pathways^9^ associated with the lists of transcriptionally up-regulated genes (orange triangles pointing upwards) and transcriptionally down-regulated genes (blue triangles pointing downwards), plotted according to the false discovery rate (FDR)-corrected *p* value of the enrichment. Full results, including pathway IDs, are in Supplementary Table 3. **(C)** Volcano plot showing the relative change in abundance of cellular transcripts and the FDR-corrected *p* value for differential expression between the mock and infected samples (n=2 biological replicates). Grey vertical lines indicate a transcript abundance fold change of 2. Genes which have fold changes greater than this threshold and a *p* ≤ 0.05 value of less than 0.05 are considered significantly differentially expressed and coloured orange if up-regulated and blue if down-regulated. Selected genes are annotated in red. **(D)** Volcano plot showing the relative change in translation efficiency of cellular transcripts, and the FDR-corrected *p* value, between the mock and infected samples (n=2 biological replicates). Colours and fold change and *p* value thresholds as in C. **(E)** Analysis of RPFs mapping to *Atf4* (NCBI RefSeq mRNA NM_009716). Cells were infected with MHV-A59 or mock-infected and harvested at 5 h p.i. (libraries from replicate 2) or 8 h p.i. RPFs are plotted at the inferred position of the ribosomal P site and coloured according to the phase of translation: pink for 0, blue for +1, yellow for +2. The main ORF (0 frame) is shown at the top in pink, with start and stop codons in all three frames marked by green and red bars (respectively) in the three panels below. The two yellow rectangles are in the +2 frame indicate the known *Atf4* uORFs (the first of which is only three codons in length). Dotted lines serve as markers for the start and end of the features in their matching colour. Note that read densities are plotted as reads per million host-mRNA-mapping reads, and that bar widths were increased to 12 nt to aid visibility, and therefore overlap, and were plotted sequentially starting from the 5′ end of the transcript. **(F)** Plot of log_2_(fold changes) of translation efficiency (TE) vs transcript abundance for all genes included in both analyses. Grey lines indicate fold changes of 2. Fold changes are plotted without filtering for significant *p* values. Selected genes are marked: genes up-regulated predominantly by one of either transcription or TE are marked in orange (upper middle and right middle sections), and *Chop*, which is up-regulated at the level of both transcription and TE, is marked in green (top right section).

Here, we characterise global changes in the host translatome and transcriptome during murine coronavirus (MHV) infection using RNA sequencing (RNASeq) and ribosome profiling (RiboSeq). MHV is a member of the *Betacoronaviru*s genus and is widely used as a model to study the replication and biology of members of the genus. In this analysis, the UPR is one of the most significantly enriched pathways. We further confirm the activation of all three branches of the UPR in MHV-infected cells. Extending our investigation to SARS-CoV-2, we find that infection with this novel CoV also activates all three UPR pathways. Moreover, we demonstrate that individual over-expression of SARS-CoV-2 ORF8 and S proteins is sufficient to induce the UPR. Remarkably, pharmacological inhibition of the UPR had a dramatic negative effect on MHV and SARS-CoV-2 replication, suggesting that CoVs may subvert the UPR to their own advantage. These results reveal that pharmacological manipulation of the UPR can be used as a therapeutic strategy against coronavirus infection.

## Results

### Differential gene expression analysis of murine cells infected with MHV-A59

To survey genome-wide changes in host transcription and translation during CoV infection, murine 17 clone 1 cells (17 Cl-1) were infected with recombinant MHV-A59 at a multiplicity of infection (MOI) of 10, or mock-infected, in duplicate and harvested at 5 hours post-infection (h p.i.). Lysates were subjected to RNASeq and parallel RiboSeq [6, 7], which allows global monitoring of cellular translation by mapping the positions and abundance of translating ribosomes on the transcriptome with sub-codon precision. Quality control analysis confirmed the libraries were of high quality (S1 Figure, S1 Table).

To assess the effects of MHV infection on cellular transcript abundance, differential expression analysis was performed at 5 h p.i. with DESeq2 [8] (Fig 1B and C, S2 and S3 Tables). At this timepoint, viral RNA synthesis approaches a maximum, but it precedes the onset of cytopathic effects such as syncytium formation [7]. Between infected and mock-infected conditions, genes with a fold change ≥2 and a false discovery rate (FDR)-corrected *p* value of ≤0.05 were considered to be significantly differentially transcribed (S2 Table). To determine the biological pathways involved in the response to infection, we carried out Reactome pathway enrichment analysis [9] on the lists of significantly differentially transcribed genes (Fig 1B, S3 Table). The most significantly enriched pathway associated with transcriptionally up-regulated genes was “Unfolded Protein Response” (R-HSA-381119, *p* = 1.1×10^−10^), and pathways denoting the three branches of the UPR (ATF6 branch: R-HSA-381183, PERK branch: R-HSA-380994, IRE1α branch: R-HSA-381070) were also significantly enriched (S3 Table). Consistent with this, gene ontology (GO) term enrichment analysis of the transcriptionally up-regulated gene list revealed that UPR-related GO terms, such as “response to unfolded protein” (GO:0006986), were significantly enriched (S3 Table). Many of the enriched pathways and GO terms associated with transcriptionally down-regulated genes are related to protein synthesis, again highlighting this as a key theme of the host response.

We provide the full database of differentially expressed genes and enriched pathways/GO terms for further exploration (S2 and S3 Tables) but in this manuscript we will focus predominantly on the UPR, which has been recognised as a host response to several CoVs due to the extensive dependence of CoV replication on the ER [4]. Accordingly, some of the most differentially transcribed genes are involved in the UPR, such as *Herp* (also known as *Herpud*), *Chac1*, *Bip* (also known as *Grp78* or *Hspa5*), *Chop* (also known as *Ddit3* or *Gadd153*) and *Grp94* (also known as *Hsp90b1*) (Fig 1C).

To evaluate differences at the level of translation, we calculated relative translation efficiencies (TE; defined herein as the ratio of ribosome-protected-fragment [RPF] to total RNA density in the CDS of a given gene) at 5 h p.i. using Xtail [10], applying the same fold change and *p*-value thresholds as for the transcription analysis. As shown in Fig 1D, several of the translationally up-regulated genes encode key proteins involved in activation of the UPR, for example ATF4, ATF5 and CHOP, which are effector transcription factors [11–16]. GADD34 (also known as MYD116/PPP1R15A), a protein that acts as a negative regulator to diminish prolonged UPR activation [17, 18], was also translationally up-regulated.

Given that UPR activation can lead to eIF2α phosphorylation and host translational shut-off, we investigated whether the list of mRNAs found to be preferentially translated during MHV infection was enriched for genes resistant to translational repression by phosphorylated eIF2α (p-eIF2α) (Materials and Methods, S4 Table). We found a 9.15-fold enrichment of p-eIF2α resistant genes (*p* = 1.42×10^−4^, Fisher Exact Test). Resistance to the effects of p-eIF2α has been linked to the presence of efficiently translated upstream open reading frames (uORFs) in the 5’ UTR [11–16,19]. To investigate this in our dataset, we analysed ribosome occupancy of the main ORF compared to the uORFs on *Atf4*, a well-studied example(12) (Fig 1E). Translation of the short (three codon) uORF1 was observed under all conditions. In mock-infected samples, uORF2 was efficiently translated, largely precluding translation of the main ORF (pink). In contrast, in MHV-infected cells, a large proportion of ribosomes scan past uORF2 to translate the main ORF. This is consistent with previous studies on *Atf4* translation under conditions of eIF2α phosphorylation, in which many ribosomes cannot reassemble a competent initiation complex before reaching uORF2 [11, 12]. This facilitates increased production of Atf4 even when translation of most mRNAs is inhibited.

Comparison of the fold changes at the transcriptional and translational level for individual cellular mRNAs provides insight into the overall effect on gene expression (Fig 1F). Genes regulated in opposing directions transcriptionally and translationally likely result in a small overall change in expression, whereas genes regulated only in one direction likely result in a greater overall change. Many UPR genes fall into the latter category (orange points, top-centre and right-centre), reflecting published knowledge about the induction of these genes specifically at the transcriptional [20–23] or translational level [12–16,19]. *Chop* (green point, upper-right) is a rare example of a gene that is significantly up-regulated both transcriptionally and translationally during MHV infection. This reflects the fact that it is transcriptionally induced by ATF4 during UPR activation and translationally p-eIF2α-resistant [24, 25].

Together, the ribosome profiling results highlight the UPR as a key pathway in the host response to MHV infection, with many of the greatest expression changes observed for UPR-related genes.

### MHV infection and activation of the unfolded protein response

To further explore the extent of UPR activation during MHV infection, we investigated each of the three branches individually (Fig 1A), building on the work of several groups [26–30].

#### Monitoring the PERK-eIF2α-ATF4 branch

Upon ER stress, PERK oligomerises and auto-phosphorylates [31]. Activated PERK phosphorylates the α-subunit of eIF2 which in turn impairs recycling of inactive eIF2-GDP to active eIF2-GTP, resulting in a general shutdown of protein synthesis [32]. However, translation of ATF4 is increased in this situation [12,33,34] leading to the induction of its target genes *Chop* and *Gadd34* (Fig 1A, right). To assay PERK activation, we monitored expression of PERK, CHOP, ATF4 and p-eIF2α, by qRT-PCR and western blotting. 17 Cl-1 cells were infected with MHV-A59 or incubated with tunicamycin and harvested at 2.5, 5, 8 and 10 h. Tunicamycin, used as a positive control, is a pharmacological inducer of ER stress which activates all UPR signalling pathways. From 5 h p.i. onwards in MHV-infected cells, and at all timepoints in tunicamycin-treated cells, ATF4 and p-eIF2α were detected and multiple bands were observed for PERK (Fig 2A) corresponding to the auto-phosphorylated species, indicative of activation of this kinase upon ER stress. In addition, as shown in Fig 2B, *Chop* and *Gadd34* mRNA levels in MHV-infected cells (blue squares) increased from 2.5 to 8 h p.i., similarly to tunicamycin-treated cells (red circles), indicating their induction by the transcription factor ATF4.

**Figure 2:**
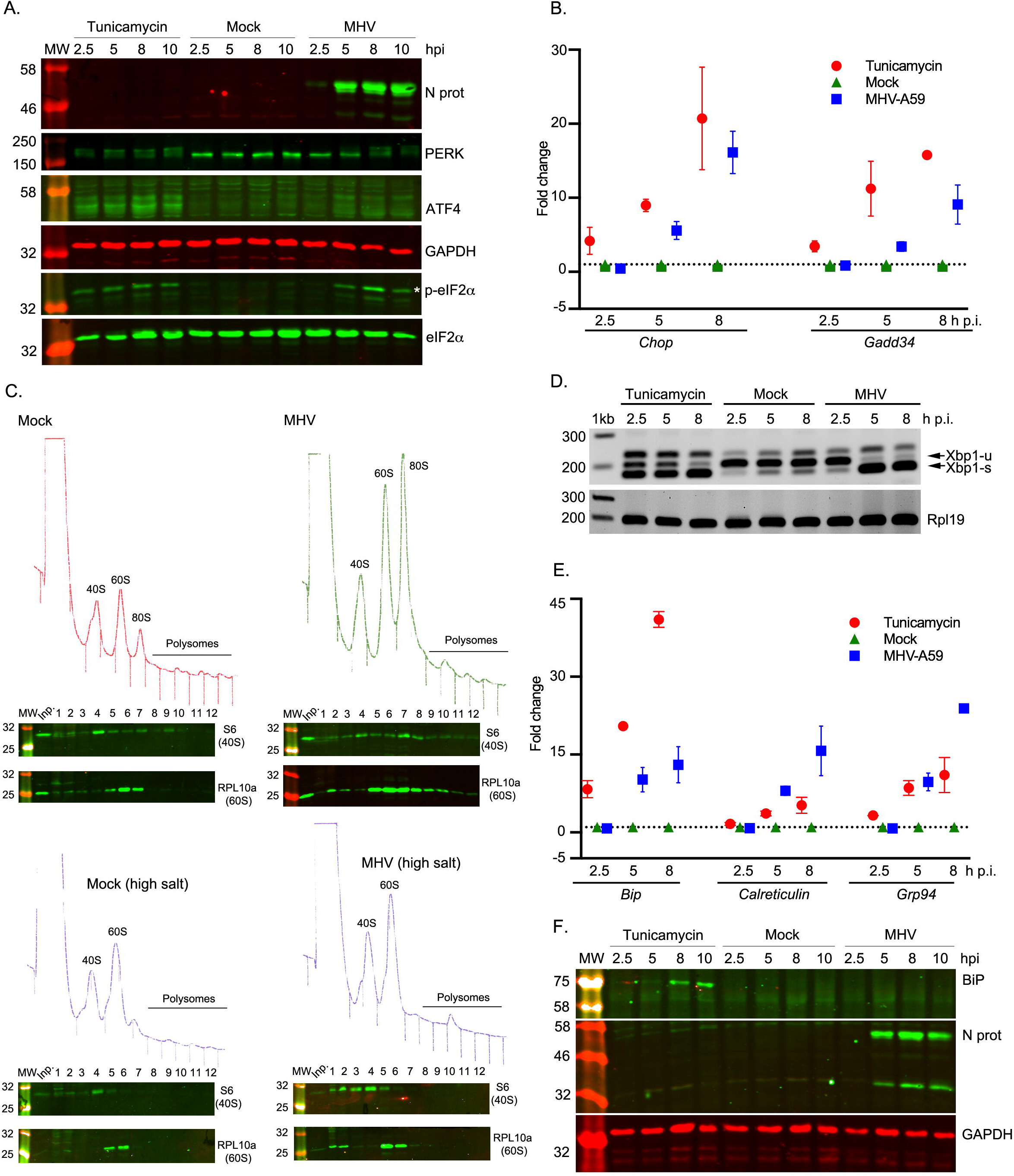
MHV infection and activation of the unfolded protein response. 17 Cl-1 cells were incubated in the presence of tunicamycin (2 µg/ml) or infected with MHV-A59 (MOI 5) and harvested at 2.5, 5, 8 and 10 h p.i. **(A)** Cell lysates were separated by 12% SDS-PAGE and immunoblotted using anti-ATF4, anti-p-eIF2α, anti-eIF2α, anti-PERK and anti-N antibodies (green fluorescent secondary antibody). GAPDH was used as a loading control (red fluorescent secondary antibody). Molecular masses (kDa) are indicated on the left and the p-eIF2α band is indicated by a white asterisk. **(B)** RT-qPCR of *Chop* and *Gadd34* mRNA for three biological replicates of a timecourse of MHV infection or tunicamycin treatment. Data are normalised using *Rpl19* as a housekeeping gene and presented as fold change of expression relative to mock-infected cells (marked as a dashed line). **(C)** Mock-infected (left upper panel) and MHV-infected (right upper panel) 17 Cl-1 cells were harvested at 5 h p.i. Cytoplasmic lysates were resolved on 10–50% sucrose density gradients. Gradients were fractionated and fractions monitored by absorbance (A_254_ nm). Twelve numbered fractions were collected and proteins extracted, resolved by 12% SDS-PAGE and analysed by immunoblotting using the indicated antibodies (anti-S6 as 40S marker, anti-RPL10 as 60S marker). Mock-infected (left lower panel) and MHV-infected (right lower panel) 17 Cl-1 cells were harvested at 5 h p.i. in high-salt lysis buffer (400 mM KCl) and analysed as described above. Molecular masses (kDa) are indicated on the left. Lane “Inp” contains whole cell lysate. **(D)** RT-PCR analysis of *Xbp1-u* and *Xbp1-s* mRNAs. Total RNA (0.5µg) was subjected to RT-PCR analysis using primers flanking the *Xbp1* splice site. PCR products were resolved in a 3% TBE-agarose gel and visualised by ethidium bromide staining. *Rpl19* RT-PCR product was used as a loading control. Molecular size markers (nt) are indicated on the left. **(E)** RT-qPCR of *Bip*, *calreticulin* and *Grp94* mRNA for three biological replicates of a timecourse of MHV infection or tunicamycin treatment. Data are normalised as in B. (**F**) Cell lysates were analysed by 12% SDS-PAGE and immunoblotted using anti-BiP and anti-N antibodies. GAPDH was used as a loading control.

Virus-induced inhibition of translation as a consequence of eIF2α phosphorylation was confirmed by analytical polysome profiling in 17 Cl-1 cells (Fig 2C, upper panel), revealing the accumulation of monosomes (80S) in MHV-infected cells at 5 h p.i. In higher salt profiles (400 mM KCl; Fig 2C, lower panel), where 80S ribosomes lacking mRNA dissociate into constituent subunits, a large reduction in 80S ribosomes was seen. These data are highly consistent with inhibition of translation initiation and show that the vast majority of 80S ribosomes accumulating at this time point are not mRNA-associated. These data support the view that MHV infection leads to translational shut-off via inhibited initiation, consistent with the effects of eIF2α phosphorylation.

#### Monitoring the IRE1α-XBP1 branch

Activated IRE1α (Fig 1A, left) removes a 26-nt intron from unspliced *Xbp1* (*Xbp1-u*) mRNA leading to a translational reading frame shift and a longer protein [23, 35]. The product of spliced *Xbp1* mRNA (XBP1-s) is an active transcription factor that up-regulates the expression of ER-associated degradation (ERAD) components and ER chaperones. To study this, we analysed *Xbp1-u* and *Xbp1-s* mRNAs by reverse transcriptase PCR (RT-PCR), using specific primers flanking the splice site (Fig 2D). At all timepoints, *Xbp1-u* was the predominant form in mock-infected cells whereas *Xbp1-s* was the major species in tunicamycin-treated cells. In virus-infected cells, *Xbp1-s* became predominant at 5 h p.i. This was corroborated at the translational level in the ribosome profiling datasets, in which infected samples showed increased translation of the extended ORF (yellow) generated by splicing (S2 Figure). An increase in active XBP1-s transcription factor was further supported by the finding that two of its target genes are transcriptionally up-regulated in infected cells (*ERdj4* – 2.44-fold increase *p*=6.63×10^−08^; and *P58ipk* – 1.94-fold increase *p*=3.97×10^−11^) (S2 Table). These data indicate that the IRE1α-Xbp1 pathway is activated by MHV infection.

#### Monitoring the ATF6 branch

The ATF6 branch is activated when ATF6 translocates from the ER to the Golgi apparatus, where it is cleaved [36]. After cleavage, the amino-terminus of ATF6 (ATF6-Nt) translocates to the nucleus to up-regulate ER chaperones (Fig 1A, middle). To monitor this pathway, 17 Cl-1 cells were infected with MHV-A59 or incubated with tunicamycin and analysed by western blotting (to detect ATF6 cleavage) or by immunofluorescence (to detect ATF6 nuclear translocation) (S3A, S3B and S3C Figures). However, we were unable to detect the trimmed version of ATF6 nor a clear nuclear translocation. As ATF6-Nt was also not visible in the positive control tunicamycin-treated cells, it is likely that the antibodies used do not efficiently recognise mouse ATF6-Nt in this context.

As an alternative approach, we monitored the induction of *BiP*, *Grp94* and *calreticulin*, transcriptionally up-regulated genes in the Reactome category “ATF6 (ATF6-alpha) activates chaperone genes” (S3 Table) and known to be induced by ATF6-Nt [37, 38]. BiP mRNA or protein levels are often used as a proxy for activation of the ATF6 pathway; however, its transcription can eventually be regulated by other UPR factors such as XBP1 [39] and ATF4 [40], so it can also be used as general readout of ER stress [26, 37]. Cells were harvested at 2.5, 5 and 8 h p.i. and analysed by qRT-PCR (Fig 2E). An increase in *Bip* transcription was observed in tunicamycin-treated (red circles) and to a lesser extent in MHV-infected cells (blue squares) from 2.5 to 8 h p.i., whereas mock-infected cells (green triangles) showed no induction. Despite the transcriptional up-regulation and a noticeable increase in RiboSeq reads mapping to BiP (S3D Figure), the protein was not detectable by western blot in MHV-infected cells (Fig 2F). It is not yet clear why this is the case, although down-regulation of BiP at the protein level has previously been observed during infection with other members of the order *Nidovirales* [30, 41]. Nevertheless, an increase in *calreticulin* and *Grp94* transcription (Fig 2E) was observed in tunicamycin-treated cells (red circles) and to a greater extent in MHV-infected cells (blue squares) especially at 8 h p.i. This indicates that the ATF6 pathway is highly up-regulated during MHV-infection. Together with our studies of PERK-eIF2α-ATF4 and IRE1α-Xbp1 above, these data confirm that MHV infection induces all three branches of the UPR.

### Effect of UPR inhibitors on MHV replication

Based on the strong UPR activation brought about by MHV infection, we hypothesised that pharmacological manipulation of this pathway could be used to modulate viral replication. First, we determined cell viability after drug treatment using Cell Titre Glo and trypan blue exclusion assays (S4 Figure). Subsequently, we evaluated the inhibitory effect of four different UPR inhibitors (UPRi) on each one of the UPR branches in cells infected with MHV for 8 h at MOI 5 (S5 Figure).

GSK-2606414 (henceforth referred to as PERKi) is a specific inhibitor of PERK [42, 43]. As expected, PERKi treatment prevented autophosphorylation of PERK and reduced phosphorylation of its substrate, eIF2α (S5A Figure), effectively blocking this branch of the UPR. Pulse labelling of infected cells for one hour at 5 h p.i. revealed a modest increase of both viral and host protein synthesis, with no effect on mock-infected cells (S5B Figure). Analytical polysome profiling of MHV-infected cells treated with 5 *μ*M PERKi for 5 h (S5C Figure) revealed a decrease in the accumulation of monosomes (80S) compared to MHV-infected cells at 5 h p.i. (Fig 2C, upper right panel), indicating a relief of translation inhibition.

Integrated stress response inhibitor (ISRIB) acts downstream of eIF2α in the PERK pathway by preventing p-eIF2α from binding and inhibiting eIF2B [44]. Therefore, eIF2B can recycle eIF2-GDP to active eIF2-GTP, and translation initiation can still occur, despite the levels of p-eIF2α remaining unchanged. Inhibition of the PERK pathway downstream of eIF2α is evident from the decrease in *Chop* transcription in MHV-infected cells treated with 2 µM ISRIB (S5D Figure).

STF-083010 (henceforth referred to as IREi) is a specific IRE1α endonuclease inhibitor that does not affect its kinase activity [45]. In MHV-infected cells treated with IREi at 60 *μ*M (8 h p.i., S5E Figure) the unspliced form of Xbp1 was more prominent compared to the untreated MHV-infected cells, indicating a block in the endonuclease activity of this enzyme.

AEBSF, a serine protease inhibitor, prevents ER stress-induced cleavage of ATF6 resulting in inhibition of transcriptional induction of ATF6 target genes [46]. We investigated the induction of ATF6 target genes in MHV-infected cells treated with 100 *μ*M AEBSF as previously described. As anticipated, *calreticulin* and *Grp94* transcription was greatly reduced in AEBSF-treated cells (S5F Figure).

Having shown these compounds effectively inhibit the UPR in the context of infection, we moved on to assess whether this could lead to an inhibition of viral replication. Cells were infected with MHV at MOI 5 and treated with the UPRi. At 8 h p.i., tissue culture supernatant was harvested and released progeny quantified by plaque assay. We found significant reductions in virus titres for all UPRi treatments in comparison to control cells, with fold reductions of between ∼two-fold (IREi) and ∼six-fold (ISRIB) (Fig 3A). This supports our hypothesis that modulation of the UPR can have antiviral effects.

**Figure 3:**
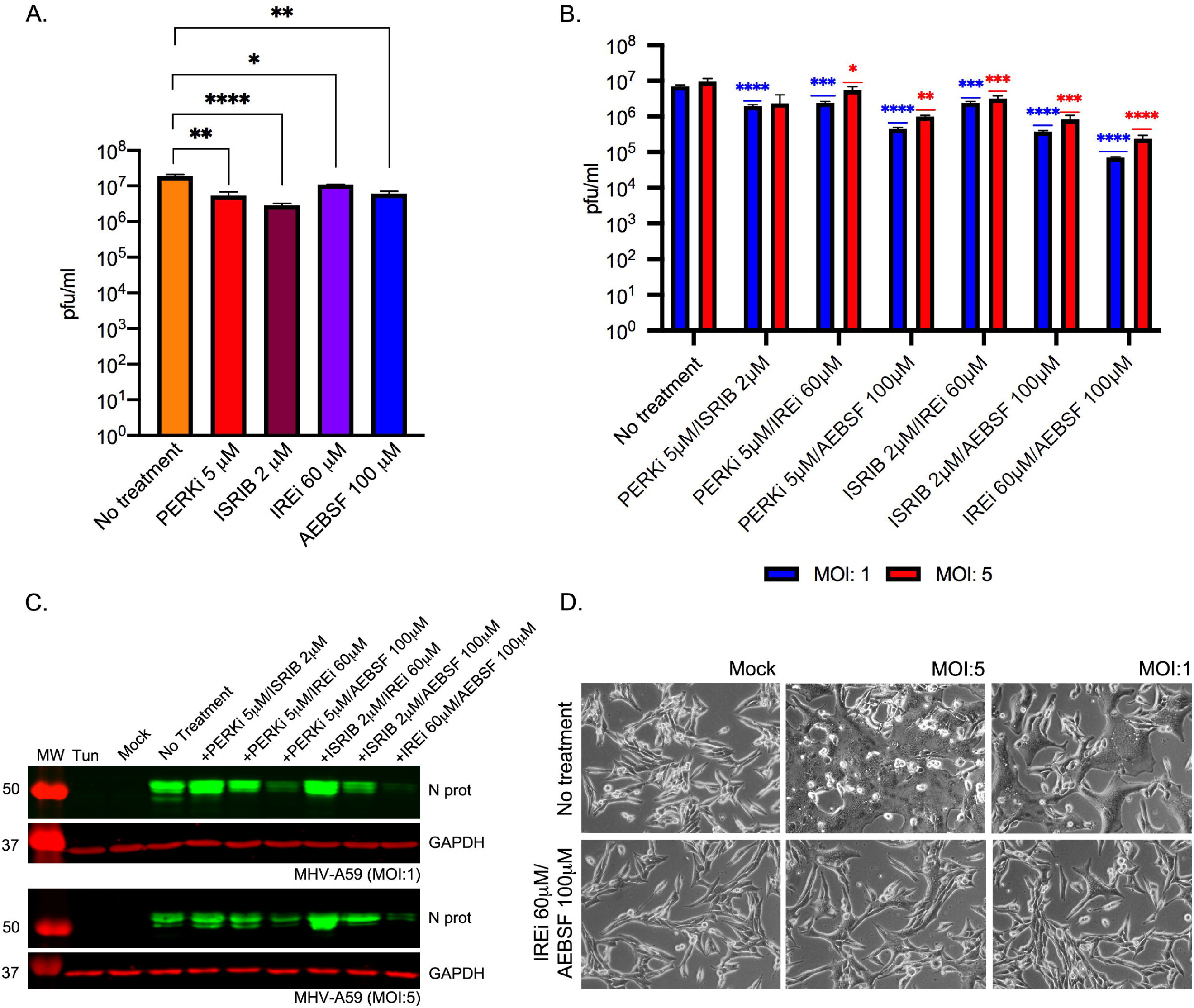
Effect of UPR inhibitors on MHV replication. (**A**) MHV-infected cells (MOI 5) were treated with UPR inhibitors (5 *μ*M PERKi, 2 *μ*M ISRIB, 60 *μ*M IREi, or 100 *μ*M AEBSF). The inhibitors were added to the cells immediately after the virus adsorption period and maintained in the medium until cells were harvested 8 h later. Plaque assays were performed with serial dilutions of the supernatant containing released virions from 17 Cl-1 cells infected with MHV-A59 in the presence or absence of the UPR inhibitors. Values show the mean averages of the titration of three biological replicates. Error bars represent standard errors. **(B-D)** MHV-infected cells (MOI 1 and MOI 5) were treated with dual combinations of the UPR inhibitors. The inhibitors were added to the cells immediately after the virus adsorption period and maintained in the medium until cells were harvested 8 h later. (**B**) Released virions were quantified as described in A. (**C**) Cell lysates were separated by 12% SDS-PAGE and immunoblotted using anti-N and anti-GAPDH antibodies as described in Fig 2A. (**D**) Representative images of mock- and MHV-infected cells at 8 h p.i. under no-drug or IREi 60 µM/AEBSF 100*μ*M treatment conditions. All *t*-tests were two-tailed and did not assume equal variance for the two populations being compared (\**p* < 0.05, ** *p* < 0.01, *** *p* < 0.001, **** *p* < 0.0001). All *p*-values are from comparisons with the respective no-treatment control at the same MOI.

Next, we investigated whether using the UPRi in combination would have a cumulative effect on virus release. We confirmed that combination treatment conditions led to reversal of the three branches of the UPR, assayed as described above (S6 Figure). Fig 3B shows virus titres from infected cells (8 h p.i.) at MOI 1 (blue) and MOI 5 (red), treated with different UPRi combinations. Reductions in virus titre ranged from ∼four-fold, in cells incubated with PERKi and ISRIB (both targeting the PERK-eIF2α-ATF4 branch), to ∼40- and ∼100-fold (MOI 5 and 1 respectively), in cells treated with IREi and AEBSF (targeting the IRE1α and the ATF6 pathways). This was further confirmed by western blotting, demonstrating a striking decrease in N protein levels for treatment combinations where virus titres were lowest (Fig 3C). In addition, cell monolayers infected with MHV in the presence of IREi and AEBSF showed delayed cytopathic effect, as indicated by reduced syncytium formation, likely due to lower virus production (Fig 3D).

### Mechanistic analysis of the UPR activation by SARS-CoV-2 proteins

Having established the use of UPRi as a potential antiviral strategy, we moved on to study UPR activation by SARS-CoV-2, initially assaying the cellular response to individual virus proteins in the context of transfection.

Previous studies have indicated that expression of the SARS-CoV spike (S) protein can activate the PERK-eIF2α-ATF4 branch [47] whereas the MHV S protein activates the IRE1α-XBP1 pathway (28). In addition, SARS-CoV ORF3a and ORF8 were found to activate the PERK-eIF2α-ATF4 and ATF6 pathways, respectively [48, 49]. To define the UPR activation associated with the counterpart proteins from SARS-CoV-2, we expressed C-terminally-tagged S (S-HA), ORF3a (ORF3a-FLAG) and ORF8 (ORF8-FLAG) proteins in human embryonic kidney cells (HEK-293T cells). N, a structural protein which is not documented as activating the UPR, was over-expressed as a negative control (N-FLAG).

ER stress, assessed by the induction of HERP and BiP, was induced by SARS-CoV-2 S but not N (S7A Figure). The PERK-eIF2α-ATF4 branch was activated from 24 h p.t. onwards, as indicated by the phosphorylation of eIF2α and the detection of ATF4 (S7A Figure), although phosphorylation of PERK was not clearly evident. The activation of this pathway was further confirmed by the increase in *CHOP* transcription compared to mock-transfected cells (S7B Figure). The amino terminus of ATF6 (ATF6-Nt) was detected in S-transfected cells from 24 h p.t. onwards (S7A Figure), indicating activation of the ATF6 branch. Activation of the IRE1α pathway is also evident from an increase in the spliced form of *XBP1* in S protein-transfected cells (S7A Figure). Contrary to previous findings for SARS-CoV, this indicates that the expression of the SARS-CoV-2 S protein is sufficient to induce all three major signalling pathways of the UPR.

In the case of SARS-CoV-2 ORF8 transfection, IRE1α-XBP1 and ATF6 were the main pathways induced (S7C Figure), again contrasting with findings for SARS-CoV [49]. Although a slight activation of ATF4 was observed in ORF8-transfected cells at 36 h p.t. (S7C Figure), this was not accompanied by PERK nor eIF2a phosphorylation, and induction of *CHOP* transcription was lower than in S protein-transfected cells (S7B Figure). SARS-CoV-2 ORF3a transfection did not induce any of the branches of the UPR (S7C Figure).

We then asked whether the UPR induction caused by SARS-CoV-2 S and ORF8 overexpression could be reversed by treatment with UPRi. This was confirmed for each inhibitor individually (S8 Figure). Additionally, we tested this using a combination treatment condition (Fig 4), for which we selected IREi/AEBSF as this gave the most promising reduction in viral titre during MHV infection (Fig 3B). Treatment of SARS-CoV-2 S- and ORF8-transfected cells with IREi/AEBSF reduced expression of HERP and BiP to levels comparable to mock-transfected cells (Fig 4A, 36 h p.t.). This indicates the treatment successfully reversed the UPR activation by the two viral proteins. PERK pathway inhibition was evident in treated cells from the reduction in PERK and eIF2α phosphorylation (Fig 4A); however, ATF4 levels appeared to be slightly increased under these conditions, as was induction of its target gene *CHOP* (S8C Figure). ATF4 induction in the presence of IREi has been previously described [50]. Inhibition of the ATF6 and the IRE1α-XBP1 pathways was also evident, as very little ATF6-Nt and *XBP1-s* were present in IREi/AEBSF treated cells (Fig 4A and Fig 4B).

**Figure 4:**
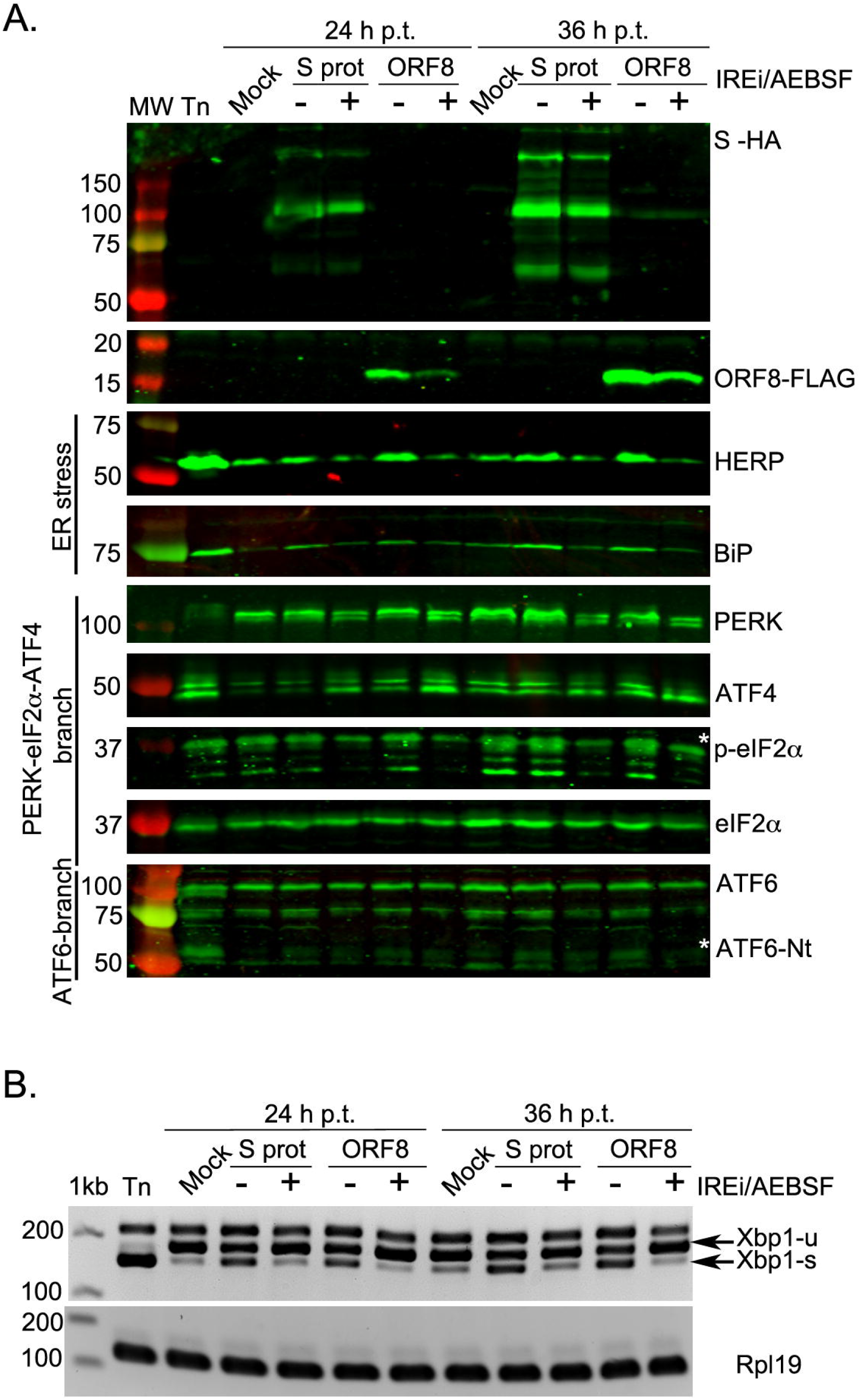
Mechanistic analysis of UPR activation by SARS-CoV-2 proteins. HEK-293T cells were transfected with plasmids encoding SARS-CoV 2 S (S-HA) or ORF8 (ORF8-FLAG), mock-transfected, or treated with tunicamycin (Tn). At 8 h p.t., cells were treated with 60 µM IREi and 100 *μ*M AEBSF and then harvested at 24 and 36 h p.t. (**A**) Cells were harvested at 24 and 36 h p.t. and cell lysates were separated by 12% SDS-PAGE and immunoblotted using anti-FLAG, anti-HA, anti-HERP, anti-BiP, anti-PERK, anti-ATF4, anti-p-eIF2α, anti-eIF2α and anti-ATF6 as described in Fig 2A. The specific p-eIF2α and ATF6-Nt bands are indicated with a white asterisk. (**B**) RT-PCR analysis of *XBP1-u* and *XBP1-s* mRNAs as described in Fig 2D. “h p.t.” = hours post-transfection.

In summary, over-expression of the S and the ORF8 proteins of SARS-CoV-2 is sufficient to activate the three branches of the UPR, and this can be reversed by UPRi treatment.

### Induction of the UPR in SARS-CoV-2-infected cells

We went on to study UPR activation in the context of SARS-CoV-2 infection. Vero CCL81 cells were infected at MOI 1 and harvested at 24 and 48 h p.i. Lysates were analysed as above. As shown in Fig 5A, the PERK-eIF2α-ATF4 branch was activated as indicated by increased phosphorylation of PERK and eIF2α. This was further confirmed by the induction of ATF4 (Fig 5A) and *CHOP* in infected cells (S9A Figure). Detection of ATF6-Nt (Fig 5A) demonstrates that the ATF6 pathway is also activated during the course of infection. In addition, activation of the IRE1α pathway was evident from an increase in the spliced form of *XBP1* in SARS-CoV-2-infected cells at 48 h p.i. (Fig 5A). We conclude that SARS-CoV-2 infection induces all three branches of the UPR.

**Figure 5:**
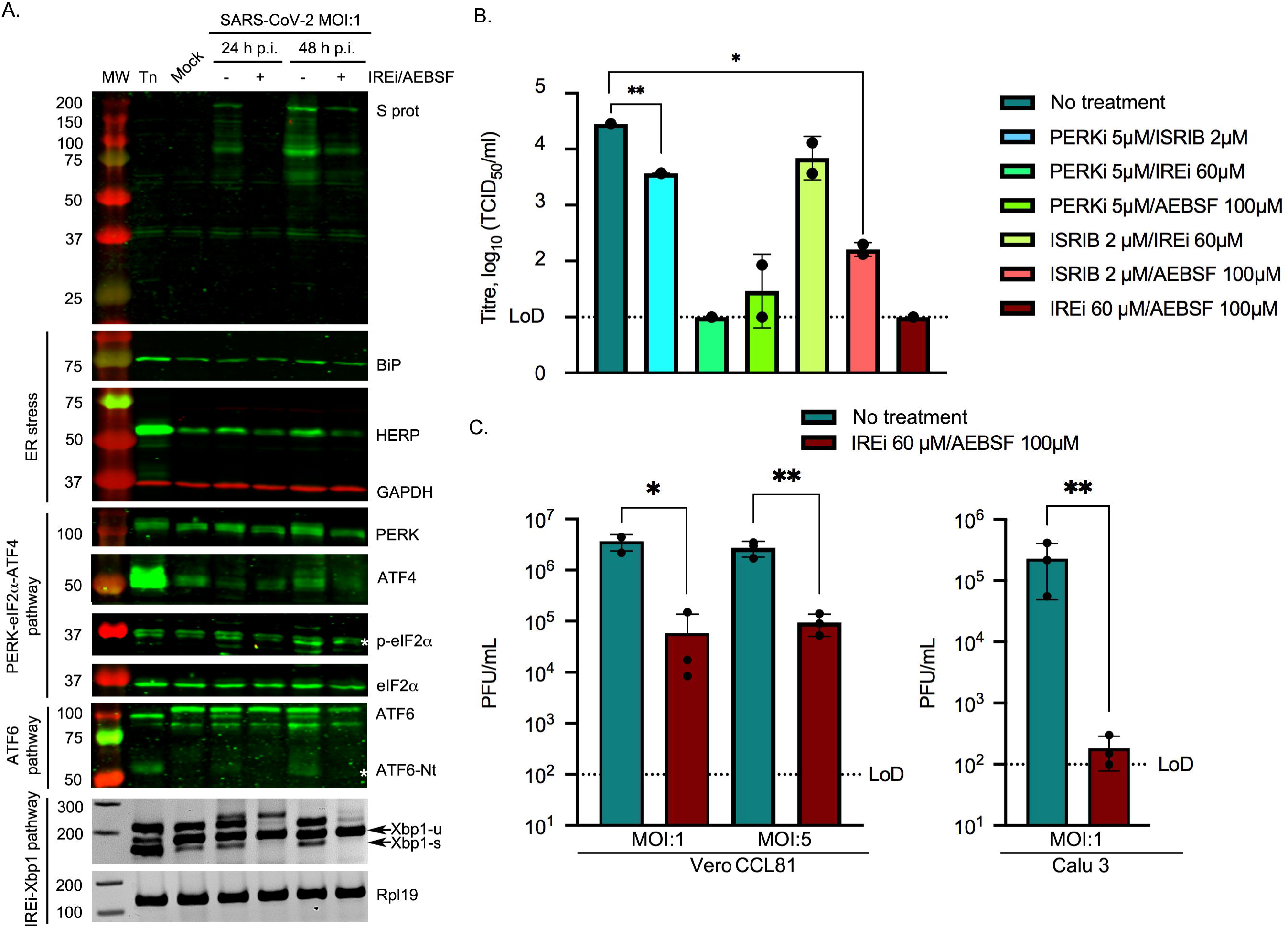
Induction of the UPR in SARS-CoV-2 infected cells and the effect of UPRi. Vero CCL81 cells were incubated in the presence of tunicamycin (2 µg/ml) or infected with SARS-CoV-2 (MOI 1) and treated with 60 µM IREi and 100 *μ*M AEBSF. The inhibitors were added to the cells immediately after the virus adsorption period and maintained in the medium until cells were harvested 24 and 48 h later. (**A**) Cell lysates (upper) were immunoblotted using anti-S, anti-BiP, anti-HERP, anti-PERK, anti-ATF4, anti-p-eIF2α, anti-eIF2α, anti-ATF6 and anti-GAPDH antibodies as described in Fig 2A. RT-PCR analysis of *Xbp1-u* and *Xbp1-s* mRNAs (lower) as described in Fig 2D. The specific p-eIF2α and ATF6-Nt bands are indicated with a white asterisk. (**B**) TCID_50_ assays were performed with serial dilutions of the supernatant containing released virions from Caco2 cells infected with SARS-CoV-2 (MOI 0.01) for 48 h in the presence or absence of the indicated UPRi combinations. (**C**) Plaque assays were performed with serial dilutions of the supernatant containing released virions from Vero CCL81 or Calu3 cells infected with SARS-CoV-2 (MOI 1 and MOI 5) for 24 h in the presence or absence of 60 µM IREi and 100 *μ*M AEBSF. Values show the mean averages of the titration of three biological replicates. Error bars represent standard errors. All *t*-tests were two-tailed and did not assume equal variance for the two populations being compared (\**p* < 0.05, ** *p* < 0.01). Replicates with titres below the limit of detection (LoD) were excluded from *t*-tests, precluding some conditions from statistical assessment.

### Effect of the IREi/AEBSF combination on SARS-CoV-2 infection

Next, we investigated whether the previously described UPRi combinations could also be used as potential antiviral drugs against SARS-CoV-2. The gastrointestinal tract is known to be one of the key sites of SARS-CoV-2 infection *in vivo* [51] so we used Caco2 cells, human intestinal cells shown to be permissive for SARS-CoV-2 infection [52, 53]. Cells were infected with SARS-CoV-2 at MOI 0.01 and treated with the different UPRi combinations. Supernatants were harvested at 48 h p.i. and released virions quantified by TCID_50_ assay (Fig 5B). Reductions in virus titre were observed and these were generally much greater than those seen for MHV (MOI 1 and 5, Fig 3B), with both the PERKi/IREi and IREi/AEBSF combinations reducing virus titres to below the limit of detection.

As the IREi/AEBSF combination had the greatest inhibitory activity against both MHV and SARS-CoV-2, we tested whether this combination could inhibit SARS-CoV-2 infection at a higher MOI. In addition to Vero CCL81 cells we employed a human lung cell line (Calu3) as a model for the primary site of SARS-CoV-2 infection, the lung [52, 53]. Cells were infected at MOI 1 or MOI 5 and virus titres assessed by plaque assays at 24 h p.i. Incubation of Vero cells with IREi/AEBSF led to a statistically significant (*p* = 0.0241 for MOI 1 and *p* = 0.0033 for MOI 5) ∼100-fold reduction in virus titre (Fig 5C, left). In Calu3 cells, IREi/AEBSF treatment had an even greater antiviral effect, reducing released virions by ∼1,000-fold (*p* = 0.0017) to at or around the limit of detection (Fig 5C, right).

Detailed analysis of the activation of the three UPR pathways under the IREi/AEBSF treatment condition was performed in SARS-CoV-2-infected Vero CCL81 cells at 24 and 48 h p.i. (Fig 5A and S9A Figure). Interestingly, in both SARS-CoV-2- and MHV-infected cells, the IREi/AEBSF combination was not only able to prevent activation of the IREi and ATF6 pathways, but also the PERK-eIF2α-ATF4 branch, as indicated by reduced phosphorylation of PERK and eIF2α (Fig 5A and S6 Figure) and reduced transcription of *Chop* (S6C Figure and S9A Figure). This may be due to the inhibition of viral replication leading to a reduced ER load, as opposed to specific inhibition of the PERK pathway. This is supported by the observation of a striking decrease in viral protein levels in infected cells treated with IREi/AEBSF (Fig 3C and Fig 5A), consistent with reduced viral replication. This reversal of CoV-induced UPR activation by the UPRi suggests that the antiviral activity of these compounds can be attributed, at least in part, to specific inhibition of the UPR, a pathway which is evidently required for efficient viral replication.

In addition to its role in UPR inhibition, AEBSF has also been reported to inhibit TMPRSS2 [54, 55], a host serine protease essential for SARS-CoV-2 cell entry [53]. To test whether AEBSF treatment inhibits SARS-CoV-2 cell entry, we transfected HEK-293T cells with TMPRSS2 and ACE2, the SARS-CoV-2 cell entry receptor [56] and incubated them with lentiviral particles pseudotyped with the SARS-CoV-2 S protein (S9B Figure). No significant inhibition of viral entry was observed upon treatment with 100*μ*M AEBSF for 4 hours, suggesting that the antiviral activity of AEBSF is predominantly due to its inhibition of the UPR.

## Discussion

This study reveals that all three branches of the UPR are activated upon MHV and SARS-CoV-2 infection, and highlights this as a very prominent pathway in the host response. The UPR was the most significantly enriched Reactome pathway associated with genes transcriptionally up-regulated during MHV infection and, consistent with previous studies, we show activation of all three branches of the UPR by MHV [26, 28]. Confirming the importance of this in SARS-CoV-2 infection, ER-related GO/KEGG terms are enriched in the differentially expressed genes lists of several proteomics/transcriptomics studies on SARS-CoV-2-infected cells [30,57–60]. This is also a very prominent theme in proteomics studies identifying host interaction partners of SARS-CoV-2 proteins, in which ER proteins are reproducibly found [59,61,62]. In one such study, “response to endoplasmic reticulum stress” was the most highly enriched biological process GO annotation associated with the host interaction partners [62]. This suggests that SARS-CoV-2, like other CoVs [63–65], enacts a finely tuned modulation of the UPR that may involve direct interactions with its components. Despite this, the activation of the three branches of the UPR by SARS-CoV-2 has not been previously described, although it has been characterised for other CoVs [4,30,63,66–70] including the closely related SARS-CoV [28,47–49,71–75]. Here we show that, like MHV, SARS-CoV-2 infection induces all three branches of the UPR, in contrast to results from SARS-CoV infection, in which only the PERK branch was activated [28,71,72].

Over-expression of the individual SARS-CoV-2 S or ORF8 proteins initiates UPR signalling. S protein was found to induce all three branches of the UPR in contrast to the counterpart protein of SARS-CoV, which appears to induce exclusively the PERK pathway [47]. Similarly, we identify ORF8 of SARS-CoV-2 as an inducer of both the IRE1α and ATF6 branches of the UPR, whereas the SARS-CoV equivalent has been shown to activate only ATF6 [49]. These differences can partly be explained by sequence divergence between the two viruses [76]. SARS-CoV-2 ORF8, for example, lacks the VLVVL motif that causes SARS-CoV ORF8 (specifically ORF8b) to aggregate and trigger intracellular stress pathways [74]. Furthermore, SARS-CoV ORF8ab was shown to mediate activation of the ATF6 pathway through a direct interaction with the ATF6 ER-lumenal domain [49], although it is undetermined whether the corresponding interaction occurs with SARS-CoV-2 ORF8. Recent proteomics-based interactome studies have identified interactions between SARS-CoV-2 ORF8 and several ER quality control proteins [59, 61], which could contribute to the ORF8-induced UPR induction observed in our study. Alterations to this key UPR modulator have important ramifications: mutation or deletion of ORF8 in naturally occurring strains of SARS-CoV and SARS-CoV-2 correlate with milder disease and, in the latter case, lower incidence of hypoxia [77–80].

Here, we also demonstrate the importance of UPR activation to CoV infection by showing that pharmacological inhibition of the UPR leads to significant reductions in titres of virions released from MHV- and SARS-CoV-2-infected cells. Simultaneous inhibition of the IRE1α and ATF6 pathways by STF-083010 and AEBSF respectively, was particularly effective, reducing virus titres by up to ∼1,000-fold. These drugs have been extensively used in preclinical studies for neurodegenerative diseases, cancer and pulmonary fibrosis [45,81–85]. Thus the STF-083010/AEBSF combination is a promising antiviral candidate to rapidly progress into a clinical trial.

To date, the development of antivirals against SARS-CoV-2 has focused on drugs targeting virus replication, such as remdesivir. However, these antiviral therapies do not take into account that the pathophysiology associated with COVID-19 is mostly related to an aberrant cellular response. In some clinical manifestations of COVID-19, an exacerbated UPR could play a key role [86–88]. For example, activation of ER stress and the UPR is one of the major triggers of endothelial dysfunction [89, 90], which is associated with acute respiratory distress syndrome (ARDS) [91], a diffuse inflammatory lung injury present in 20-67% of hospitalised patients [92, 93]. Other clinical manifestations of COVID-19 such as thromboembolism, cerebro- and cardiovascular diseases and neurological complications, are also associated with endothelial dysfunction [94]. Furthermore, a recognised sequela of COVID-19 is pulmonary fibrosis [95], which can develop in up to 17% of COVID-19 patients [96]. Pulmonary fibrosis is a severe form of interstitial lung disease characterised by progressive dyspnea, hypoxemia, and respiratory failure due to the presence of patchy areas of fibrotic tissue. ER stress and UPR activation are known to be involved in the development and progression of this fibrotic disease [97]. This suggests that UPR activation in response to SARS-CoV-2 infection contributes to the lung pathophysiology associated with COVID-19. Therefore, the UPR inhibitors used in this study could have a dual therapeutic effect, not only contributing to the reduction of viral burden in patients, but also diminishing the pathophysiology associated with COVID-19. In addition, the idea of targeting an exaggerated cellular response instead of the virus itself substantially reduces the chances of generating virus escape mutants.

## Materials and Methods

### Cells and viruses

Murine 17 clone 1 (17 Cl-1), Calu3 (ATCC, HTB-55) and Vero (ATCC, CCL81) cells were maintained in Dulbecco’s modification of Eagle’s medium supplemented with 10% (vol/vol) fetal calf serum (FCS). HEK-293T cells (ATCC, CRL-11268) were cultured in DMEM supplemented with 5% FCS. Caco2 cells were a kind gift from Dr Valeria Lulla and were maintained in DMEM supplemented with 20% FCS. All cell lines were cultured in medium containing 100 U/ml penicillin, 100 µg/ml streptomycin and 1 mM L-glutamine. Cells were incubated at 37 °C in the presence of 5% CO_2_.

Recombinant MHV strain A59 (MHV-A59) was derived as described previously [98]. Upon reaching 70–80% confluence, 17 Cl-1 cells were infected with MHV-A59 at MOI 5 as described(99,100). Vero CCL81 and Calu3 cells were infected with SARS-CoV-2 (SARS-CoV-2/human/Switzerland/ZH-UZH-IMV5/2020) at two MOIs (1 and 5) for 24 or 48 h as previously described [99, 100]. Caco2 cells were infected with SARS-CoV-2 (isolate hCoV-19/Edinburgh/2/2020, a kind gift from Dr Christine Tait-Burkhard and Dr Juergen Haas) at MOI 0.01 and incubated for 48 h in MEM containing 1% L-glutamine, 1% non-essential aminoacids, 1% penicillin/streptomycin and supplemented with 2% FBS.

### Ribosomal profiling and RNASeq data

17 Cl-1 cells were grown on 100-mm dishes to 90% confluency and infected with MHV-A59 at MOI 10. At the indicated time-points, cells were rinsed with 5 ml of ice-cold PBS, flash frozen in a dry ice/ethanol bath and lysed with 400 μl of lysis buffer [20 mM Tris-HCl pH 7.5, 150 mM NaCl, 5 mM MgCl_2_, 1 mM DTT, 1% Triton X-100, 100 μg/ml cycloheximide and 25 U/ml TURBO DNase (Life Technologies)]. The cells were scraped extensively to ensure lysis, collected and triturated ten times with a 26-G needle. Cell lysates were clarified by centrifugation at 13,000 *g* for 20 min at 4 °C. Lysates were subjected to RiboSeq and RNASeq based on previously reported protocols [7, 101]. Ribosomal RNA was removed using Ribo-Zero Gold rRNA removal kit (Illumina) and library amplicons were constructed using a small RNA cloning strategy adapted to Illumina smallRNA v2 to allow multiplexing. Amplicon libraries were deep sequenced using an Illumina NextSeq500 platform. Due to the very large amounts of vRNA produced during infection, mock samples were processed separately from infected samples to avoid contamination. RiboSeq and RNASeq sequencing data have been deposited in the ArrayExpress database (http://www.ebi.ac.uk/arrayexpress) under the accession numbers E-MTAB-8650 and E-MTAB-8651.

### Computational analysis of RiboSeq and RNASeq data

Reads were trimmed for adaptor sequences, filtered for length ≥ 25 nt, and reads mapping to *Mus musculus* rRNA (downloaded from the SILVA database [102] or MHV-A59 viral RNA (GenBank accession AY700211.1) (with up to 2 mismatches) removed, as previously described [7]. The remaining reads were aligned directly to the mouse genome (FASTA and GTF gencode release M20, GRCm38, primary assembly) (with up to 2 mismatches) using STAR (parameters: --outFilterIntronMotifs RemoveNoncanonicalUnannotated --outMultimapperOrder Random) [103]. Reads on protein-coding genes were tabulated using htseq-count (version 0.9.1), covering the whole gene for differential transcription analysis (parameters: -a 0 -m union -s yes -t gene) and just the CDS for the translation efficiency analysis (parameters: -a 0 -m intersection-strict -s yes -t CDS) [104], using the GTF file from the above Gencode release as the gene feature annotation.

Differential transcription analysis was performed using DESeq2 (version 1.18.1) [8] and translation efficiency analysis with Xtail (version 1.1.5) [10]. For each analysis, low count genes (with fewer than ten counts from all samples combined) were discarded, following which read counts were normalised by the total number of reads mapping to host mRNA for that library, using standard DESeq2 normalisation. This minimises the effect of the large amount of vRNA present in infected samples. Shrinkage of the transcriptional fold changes to reduce noise in lowly-expressed genes was applied using lfcShrink (parameter: type=’normal’).

A given gene was considered to be differentially expressed if the FDR-corrected *p* value was less than 0.05 and the fold change between the means of infected and mock replicates was greater than two. Volcano plots and transcription versus TE comparison plots were generated using R and FDR-corrected *p* values and log_2_(fold change) values from the DESeq2 and Xtail analyses. All reported *p* values are corrected for multiple testing by the Benjamini-Hochberg method. Fold changes plotted in the transcription vs TE comparison are not filtered for significant *p* values before plotting.

To plot RNASeq and RPF profiles for specific transcripts, reads were mapped to the specified transcript from the NCBI genome assembly using bowtie [105] allowing two mismatches (parameters: -v 2, --best). Coordinates for known uORFs were taken from the literature [11,12,23] and the positions of start and stop codons in all frames determined. Read density (normalised by total reads mapping to host mRNA for each library, to give reads per million mapped reads) was calculated at each nucleotide on the transcript and plotted, coloured according to phase. Read positions were offset by +12 nt so that plotted data represent the inferred position of the ribosomal P site. Bar widths were increased to 12 nt (Fig 1E) or 4 nt (Supplementary Fig 2) to aid visibility and were plotted sequentially starting from the 5’ end of the transcript.

### Gene ontology and Reactome pathway enrichment analyses

Lists of gene IDs of significantly differentially expressed genes (Supplementary Table 2) were used for GO term enrichment analysis by the PANTHER web server under the default conditions (release 20190606, GO database released 2019-02-02) [106], against a background list of all the genes that passed the threshold for inclusion in that expression analysis. For Reactome pathway enrichment (version 69) [9], the same differentially expressed gene lists were converted to their human orthologues and analysed, both using the reactome.org web server to determine which pathways are significantly over-represented (FDR-corrected *p* value ≤0.05).

### Enrichment analysis for eIF2α-phosphorylation-resistant genes

Resistance to translational repression by p-eIF2α is not an existing GO term, so a list of genes reported to be p-eIF2α-resistant was constructed based on Andreev et al., 2015 [16] and references within (excluding those from IRESite, which were not found to be p-eIF2α-resistant in their study). Mouse homologues of these genes were identified using the NCBI homologene database (Supplementary Table 4). Enrichment of genes categorised as p-eIF2α-resistant amongst the genes with significantly increased translational efficiency, compared to a background of all *Mus musculus* genes included in the TE analysis with any GO annotation, was calculated using a Fisher Exact test.

### Chemicals

GSK-2606414 was a kind gift from Dr Edward Emmott and Prof Ian Goodfellow. AEBSF, STF-083010, ISRIB and tunicamycin were purchased from Sigma-Aldrich. GSK-2606414, STF-083010, ISRIB and tunicamycin were dissolved in DMSO, whereas AEBSF was dissolved in water, to the required concentrations. In all experiments, the final concentration of DMSO did not exceed 0.4%. Cytotoxicity after treatment with single and combined UPR inhibitors was measured using the Cell Titer Blue (Promega) and trypan blue exclusion kits (Sigma), following manufacturer’s instructions.

### Antibodies

The following primary antibodies were used: mouse monoclonal antibodies against MHV N and S proteins (kind gifts of Dr Helmut Wege, University of Würzburg), mouse anti-GAPDH (IgM specific, G8795, Sigma-Aldrich), mouse anti-Flag (F3165, Sigma-Aldrich), rabbit anti-HA (3724, Cell Signaling Technology), rabbit anti-PERK (ab229912, Abcam), rabbit anti-HERPUD1 (ab150424, Abcam), rabbit anti-GRP78 (BIP, ab108613, Abcam), rabbit anti-eIF2α (9722, Cell Signaling Technology), rabbit anti-phospho-eIF2α (Ser51, 9721, Cell Signaling Technology), rabbit anti-ATF4 (10835-1-AP, Proteintech), rabbit anti-ATF6 (ab203119 and ab37149, Abcam), mouse anti-S6 (2317, Cell Signaling Technology) and rabbit RPL10a (ab174318, Abcam). Secondary antibodies used for western blotting were purchased from Licor: IRDye 800CW Donkey Anti-Mouse IgG (H+L), IRDye 800CW Donkey Anti-Rabbit IgG (H+L), IRDye 680RD Goat Anti-Mouse IgG (H+L) and IRDye 680RD Goat Anti-Mouse IgM (µ chain specific).

### Plasmids and transfections

HEK-293T cells were transiently transfected with pcDNA3.1-SARS-CoV-2-S-HA (kind gift of Dr Jerome Cattin and Prof Sean Munro, MRC-LMB, Cambridge, UK), pcDNA6-SARS-CoV-2-N-FLAG, pcDNA6-SARS-CoV-2-ORF3a-FLAG and pcDNA6-SARS-CoV-2-ORF8-FLAG plasmids (kind gifts of Prof Peihui Wang, Shandong University, China) using a commercial liposome method (TransIT-LT1, Mirus). Transfection mixtures containing plasmid DNA, serum-free medium (Opti-MEM; Gibco-BRL) and liposomes were set up as recommended by the manufacturer and added dropwise to the tissue culture growth medium. Cells were harvested at 24 and 36 h post-transfection.

### Immunoblotting

Cells were lysed in 1X Laemmli’s sample buffer. After denaturation at 98 °C for 5 minutes, proteins were separated by 12% SDS-PAGE and transferred to nitrocellulose membranes. These were blocked (5% non-fat milk powder or bovine serum albumin in PBST [137 mM NaCl, 2.7 mM KCl, 10 mM Na_2_HPO_4_, 1.5 mM KH_2_PO_4_, pH 6.7, and 0.1% Tween 20]) for 30 min at room temparature and probed with specific primary antibodies at 4°C overnight. Membranes were incubated in the dark with IRDye-conjugated secondary antibodies diluted to the recommended concentrations in PBST for 1 h at room temperature. Blots were scanned using an Odyssey Infrared Imaging System (Licor).

### Analysis of *Xbp1* splicing by RT-PCR

Total RNA was isolated from infected or transfected cells as described previously [7], and cDNA synthesised from 500 ng total RNA using M-MLV Reverse Transcriptase (Promega). Mouse or human *Xbp1* and *Rpl19* were amplified using specific primers (Supplementary Table 5). Following PCR reactions, the resulting amplicons were subjected to electrophoresis in 3% agarose gels.

### Quantitative real-time PCR assays

Relative levels of mouse or human *Bip*, *Chop*, *Gadd34, Calreticulin* and *Grp94* in cDNA samples were determined by quantitative real-time PCR (qPCR) using a Rotor-Gene 3000 (Corbett Research). Reactions were performed in a final volume of 20 μl containing Hot Start Taq (1 U, QIAGEN), 3.5 mM MgCl_2_, 2.5 mM deoxynucleotides, 450 nM SYBR Green dye, 500 nM relevant forward and reverse primers (Supplementary Table 5) and 1 µl of cDNA. No-template controls were included for each primer pair, and each qPCR reaction was carried out in duplicate. Fold changes in gene expression relative to the mock were calculated by the delta delta-cycle threshold (ΔΔCt) method, and *Rpl19* was used as a normalising housekeeping gene.

### Polysome profiling

17 Cl-1 cells were infected as described above. 10 min prior to harvesting, cells were treated with cycloheximide (100 µg/ml), washed with PBS and lysed in a buffer containing 20 mM Tris HCl pH 7.5, 100 mM KCl, 5 mM MgOAc, 0.375 mM CHX, 1 mM DTT, 0.1 mM PMSF, 2 U/µl DNase I, 0.5% NP-40, supplemented with protease and phosphatase inhibitors (ThermoFisher Scientific). Following trituration with a 26-G needle (ten passes), lysates were cleared (13,000 *g* at 4 °C for 20 min) and the supernatants layered onto 12 ml sucrose density gradients (10–50% sucrose in TMK buffer: 20 mM Tris-HCl pH 7.5, 100 mM KCl, 5 mM MgCl_2_) prepared in Beckman SW41 polypropylene tubes using a Gradient Master (Biocomp). Following centrifugation (200,000 *g* for 90 min at 4 °C), fractions were prepared using an ISCO fractionator monitoring absorbance at 254 nm. Proteins were concentrated from fractions using methanol-chloroform extraction and subjected to immunoblotting analysis. Polysome profiling in higher salt conditions was carried out as described above except that the lysis buffer and sucrose density gradient contained 400 mM KCl.

### Virus plaque assays

To determine MHV-A59 titres by plaque assay, 17 Cl-1 cells in 6-well plates were infected with 400 *μ*l of 10-fold serial dilutions of sample in infection medium (Hank’s balanced salt solution containing 50 μg/ml DEAE-dextran and 0.2% bovine serum albumin - BSA). After 45 min at 37°C with regular rocking, the inoculum was removed and replaced with a 1:1 mixture of 2.4% Avicel and MEM 2X medium (20% MEM 10X, 2% non-essential aminoacids, 200 U/ml penicillin, 200 µg/ml streptomycin, 2 mM L-glutamine, 40 mM HEPES pH 6.8, 10% tryptose phosphate broth, 10% FCS and 0.01% sodium bicarbonate). Plates were incubated at 37 °C for 48 h prior to fixing with 3.7% formaldehyde in PBS. Cell monolayers were stained with 0.1% toluidine blue to visualise plaques. SARS-CoV-2 plaque assays were performed as previously described [99]). Experiments were conducted using three biological repeats.

### TCID_50_ assays

SARS-CoV-2 replication was assessed using a 50% tissue culture infective dose (TCID_50_) assay in Vero E6 cells. Supernatant derived from infected Caco2 cells was subjected to 10-fold serial dilutions. At 72 h p.i., cells were fixed and stained as previously indicated. Wells showing any sign of cytopathic effect (CPE) were scored as positive.

### Statistical analysis of virus titre results

Data were analysed in GraphPad Prism 9.0 (GraphPad software, San Diego, CA, USA). Values represent mean ± standard deviation. Statistical significance was evaluated using two-tailed *t*-tests on log_10_(virus titre) data, which did not assume equal variances for the two populations being compared, to calculate the *p*-values. Differences as compared to the control with *p* value ≤ 0.05 were considered as statistically significant, with \**p* < 0.05, ** *p* < 0.01, *** *p* < 0.001 and **** *p* < 0.0001.

## Acknowledgements

SK, ND, GD and EB would like to acknowledge Dr Holly Shelton, Dr Isabelle Dietrich and Dr Christine Reitmayer for their supervision in the CL3 suite, and Dr Christine Tait-Burkhard and Dr Juergen Haas for the SARS-CoV-2 isolate.

NI would like to thank Dr James Edgar for providing SARS-CoV-2 plasmids.

NI would like to thank Dr Luke Meredith and Prof Ian Goodfellow for providing pcDNA3.1-ACE2, pcDNA3.1-TMPRSS2, pMD2-VSV-G and pNL4.3-Luc plasmids for pseudotyped virions.

