## Supplementary Information for "Manipulation of the unfolded protein response: a pharmacological strategy against coronavirus infection"

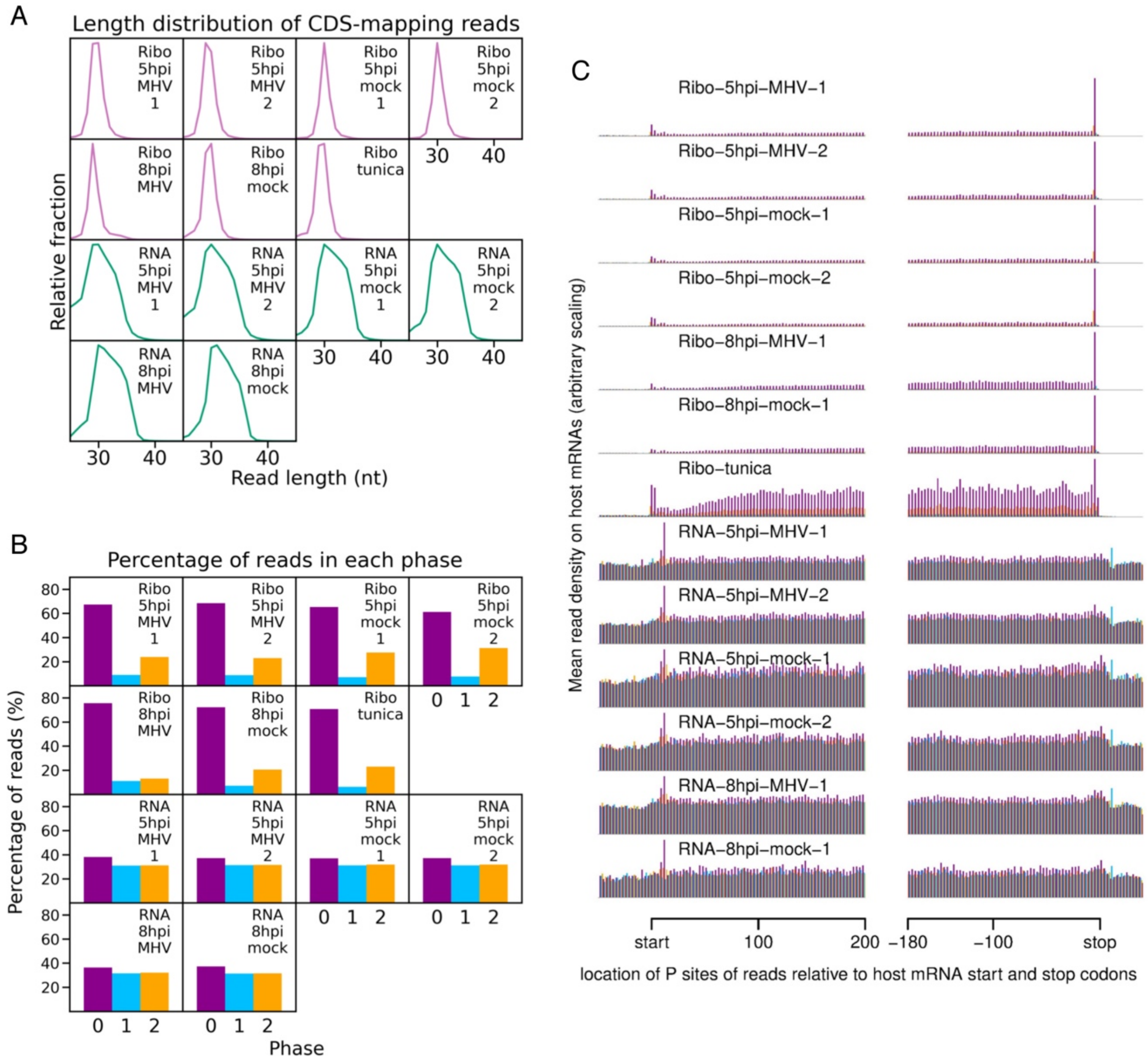


**S1 Figure. Quality control indicates sequencing data are of high quality.** **(A)** Length distribution of positive-sense reads mapping within host CDSs. For RiboSeq libraries (pink) the characteristic sharp peak at 28-29 nt is observed, reflective of the length of mRNA protected from digestion by translating ribosomes. Read lengths of RNASeq libraries (green) are determined by alkaline hydrolysis and gel purification size selection (25-34 nt), leading to a broader length distribution. (**B)** Percentage of reads (all read lengths) attributed to each phase, for positive-sense reads mapping within host CDSs. Phases correspond to which position within the codon the 5′ end of the read maps to (0: purple, 1: blue, 2: yellow). The 5′ end coordinate of RiboSeq reads is influenced by the position of the translating ribosome, leading to a clear dominance of the 0 phase. For RNASeq reads the 5′ end coordinate is determined by alkaline hydrolysis so does not result in a dominant phase. **(C)** Distribution of host mRNA-mapping reads relative to start and stop codons. Only transcripts with an annotated CDS of at least 150 codons, 5′ UTR of at least 60 nt, and 3′ UTR of at least 60 nt were included in the analysis. The total number of positive-sense reads from all these transcripts mapping to each position was plotted, with an offset of +12 relative to the 5′ end coordinate to represent the inferred ribosomal P site. Although ribosomal P site position is not relevant to RNASeq reads, these were also plotted with a +12 nt offset to facilitate comparison. Data are coloured according to phase as in B. For RiboSeq libraries there is clear triplet periodicity visible across the CDS, reflective of the length of a codon, and a large peak corresponding to terminating ribosomes - characteristic of samples harvested without drug pre-treatment. Very few RiboSeq reads map to the UTRs (and particularly the 3′ UTR), indicating very little contamination of the mRNA fraction with non-ribosome-protected-fragment reads. As expected, for RNASeq libraries the coverage does not differ greatly between the CDS and UTRs.

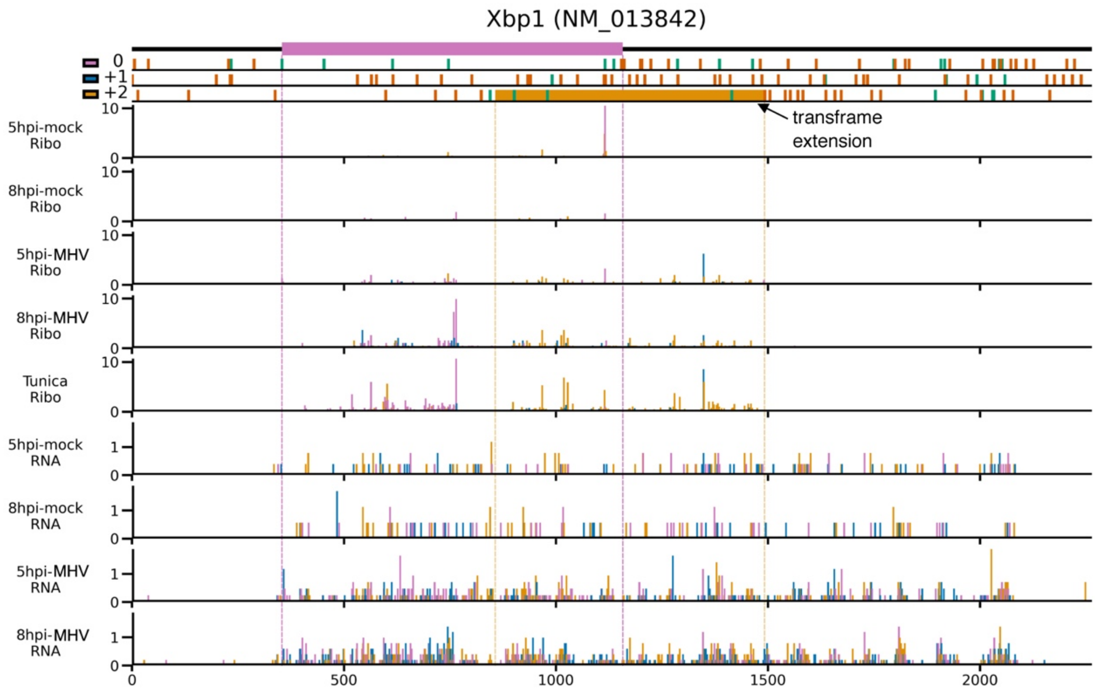


**S2 Figure. Distribution of reads mapping to specific host genes of interest.** Analysis of RPFs (mock and MHV-infected samples plus tunicamycin-treated sample) and RNASeq reads (mock and MHV-infected samples) mapping to *Xbp1-u* (NCBI RefSeq mRNA NM_013842). Cells were infected with MHV-A59 or mock-infected and harvested at 5 h p.i. or 8 h p.i. (libraries from Fig 1D and E). One sample was treated with 2 μg/ml tunicamycin, a pharmacological inducer of all three branches of the UPR, as a positive control. Reads are plotted at the inferred position of the ribosomal P site and coloured according to the phase of translation: pink for 0, blue for +1, yellow for +2. The 5′ end position of RNASeq reads is not determined by ribosome position and therefore should not show a dominant phase. The main ORF (0 frame) is shown at the top in pink, with start and stop codons in all three frames marked by green and red bars (respectively) in the three panels below. The yellow rectangle in the +2 frame indicates the extended ORF that results from splicing by IRE1. Reads resulting mainly from translation of the spliced *Xbp1-s* isoform can be seen in yellow (+2 phase), downstream of the main ORF annotated stop codon. Dotted lines serve as markers for the start and end of the features in their matching colour. Read densities are plotted as reads per million host-mRNA-mapping reads. Bar widths were increased to 4 nt to aid visibility, and therefore overlap, and were plotted sequentially starting from the 5′ end of the transcript.

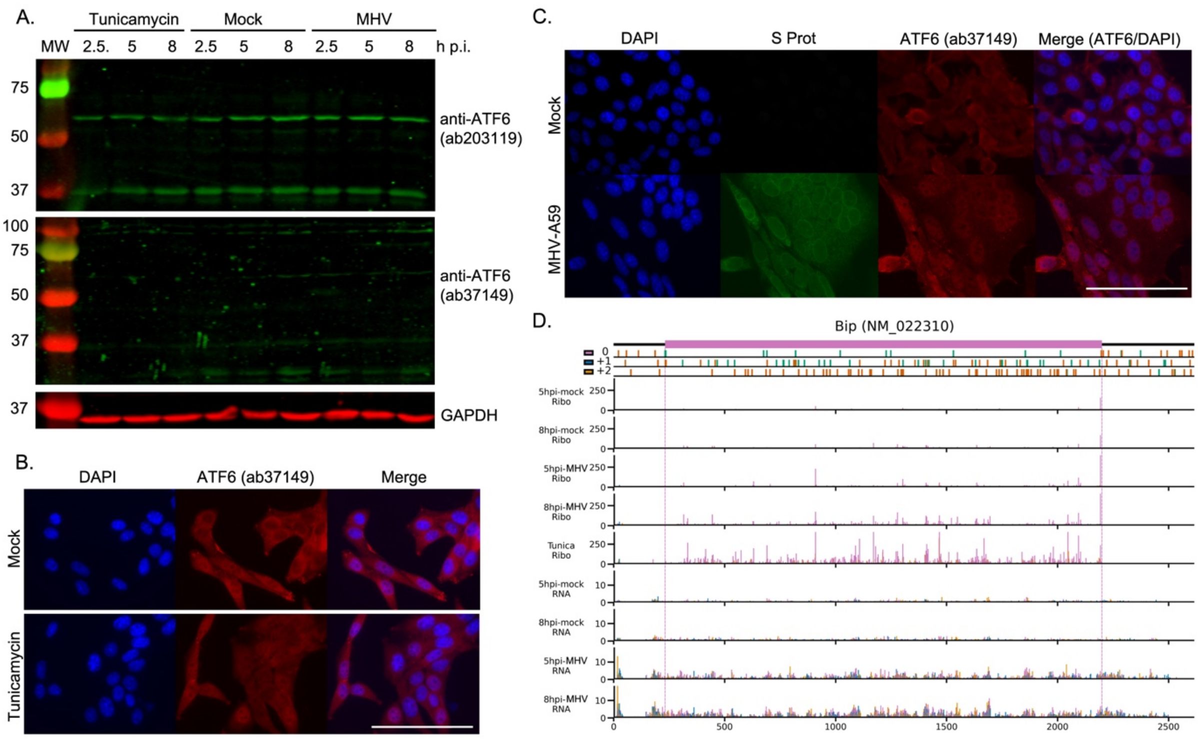


**S3 Figure. ATF6 pathway activation in MHV-infected cells.** 17 Cl-1 cells were incubated in the presence of tunicamycin (2 µg/ml) or infected with MHV-A59 (MOI 5) and harvested at 2.5, 5 and 8 h. (**A**) Cell lysates were separated by 12% SDS-PAGE and immunoblotted using anti-ATF6 (1:1000, Abcam ab203119, upper), anti-ATF6 (1:1000, Abcam ab37149, middle) and anti-GAPDH. Representative images of fixed and permeabilised cells treated with tunicamycin for 6 h (**B**) or infected with MHV for 8 h (**C**) and incubated with anti-ATF6 (red) and anti-S protein (green). Nuclei are counterstained with DAPI (blue). Images were taken in an Evos FLII microscope at 60X magnification. Scale bar: 100 µm. (**D**) Analysis of RPFs and RNASeq reads mapping to *Bip* (NM_022310). Plot constructed as described in Supplementary Fig 2. Note that in order to properly visualise RPFs across the ORF, the y-axis has been truncated at 400 reads per million host-mRNA-mapping reads for the RiboSeq samples, leaving some RPF counts for tunicamycin-treated cells and MHV-infected cells off-scale.

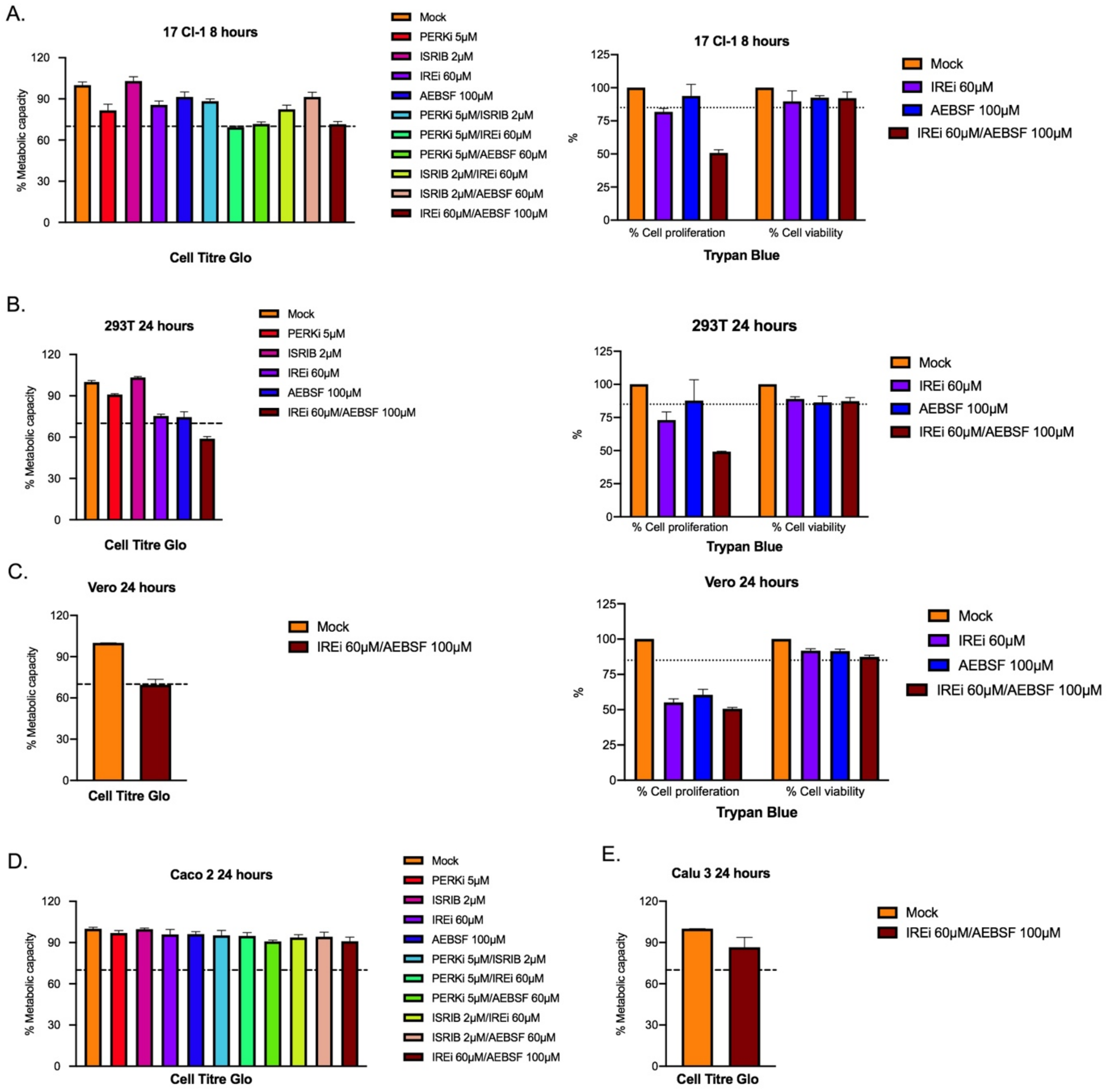


**S4 Figure. Cytotoxicity assays.** Cell viability assays of 17 Cl-1 (**A**), HEK-293T (**B**), Vero CCL81 (**C**), Caco2 (**D**) and Calu3 (**E**) cells treated with the UPR inhibitors. Experiments were performed in triplicate using CellTiter-Blue Cell Viability Assay to assess metabolic capacity and in duplicate using trypan blue exclusion assay to assess cell proliferation and viability in those cases where the metabolic capacity was ~70% (dashed line). Percentages are given relative to untreated cells. Error bars represent standard errors.

STF-083010 (IREi) has previously been described to have anti-proliferative activity [84] which is evident in highly proliferative cells such as 17 Cl-1, 293T and Vero CCL81. This property is increased in combination with AEBSF. However, cell viability in all cases tested was greater than 85% (dotted line).

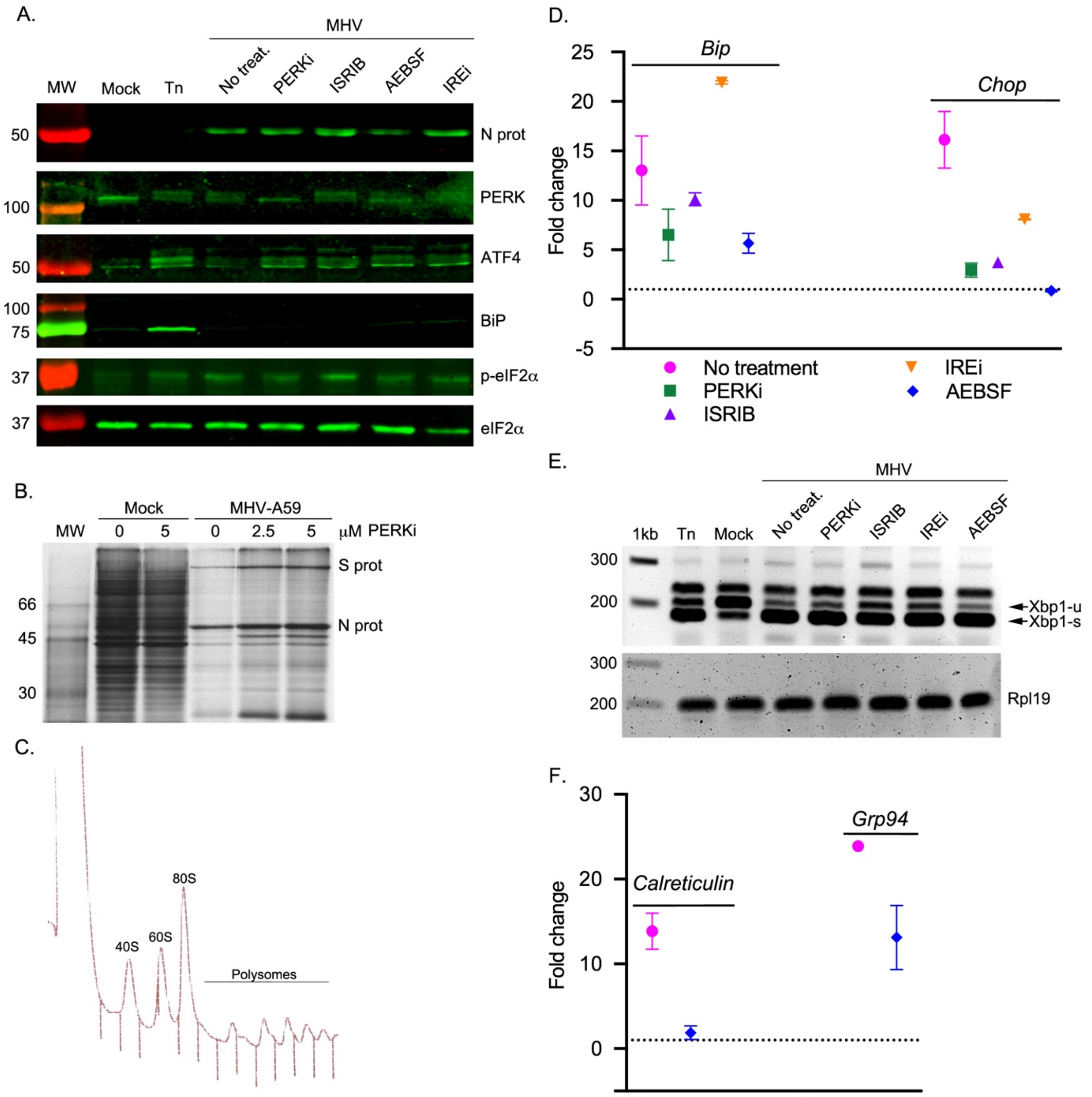


**S5 Figure. Effect of UPR inhibitors on activation of the UPR during MHV infection.** MHV-infected cells (MOI 5) were treated with UPR inhibitors (5 μM PERKi, 2 μM ISRIB, 60 μM IREi, or 100 μM AEBSF). The inhibitors were added to the cells immediately after the virus adsorption period and maintained in the medium until cells were harvested 8 h later. (**A**) Cell lysates were immunoblotted using anti-N, anti-PERK, anti-ATF4, anti-BiP, anti-p-eIF2α and anti-eIF2α antibodies as described in Fig 2A. (**B**) 17 Cl-1 cells infected with MHV-A59 and treated with 0, 2.5 or 5 μM of PERKi were metabolically pulse-labeled with [$^{35}$S]Met for 1 h at 5 h p.i. Cells were lysed just after pulse and subjected to 10% SDS-PAGE followed by autoradiography. (**C**) Polysome profiling as described in Fig

2C of MHV-infected cells at 5 h p.i. treated with 5 μM of PERKi. (**D**) RT-qPCR of *Bip* and *Chop* mRNA from two biological replicates of MHV-infected cells treated with UPR inhibitors as described in Fig 2B. (**E**) RT-PCR analysis of *Xbp1-u* and *Xbp1-s* mRNAs as described in Fig 2D. (**F**) RT-qPCR of *calreticulin* and *Grp94* mRNA from two biological replicates of MHV-infected cells treated with UPR inhibitors as described in Fig 2B.

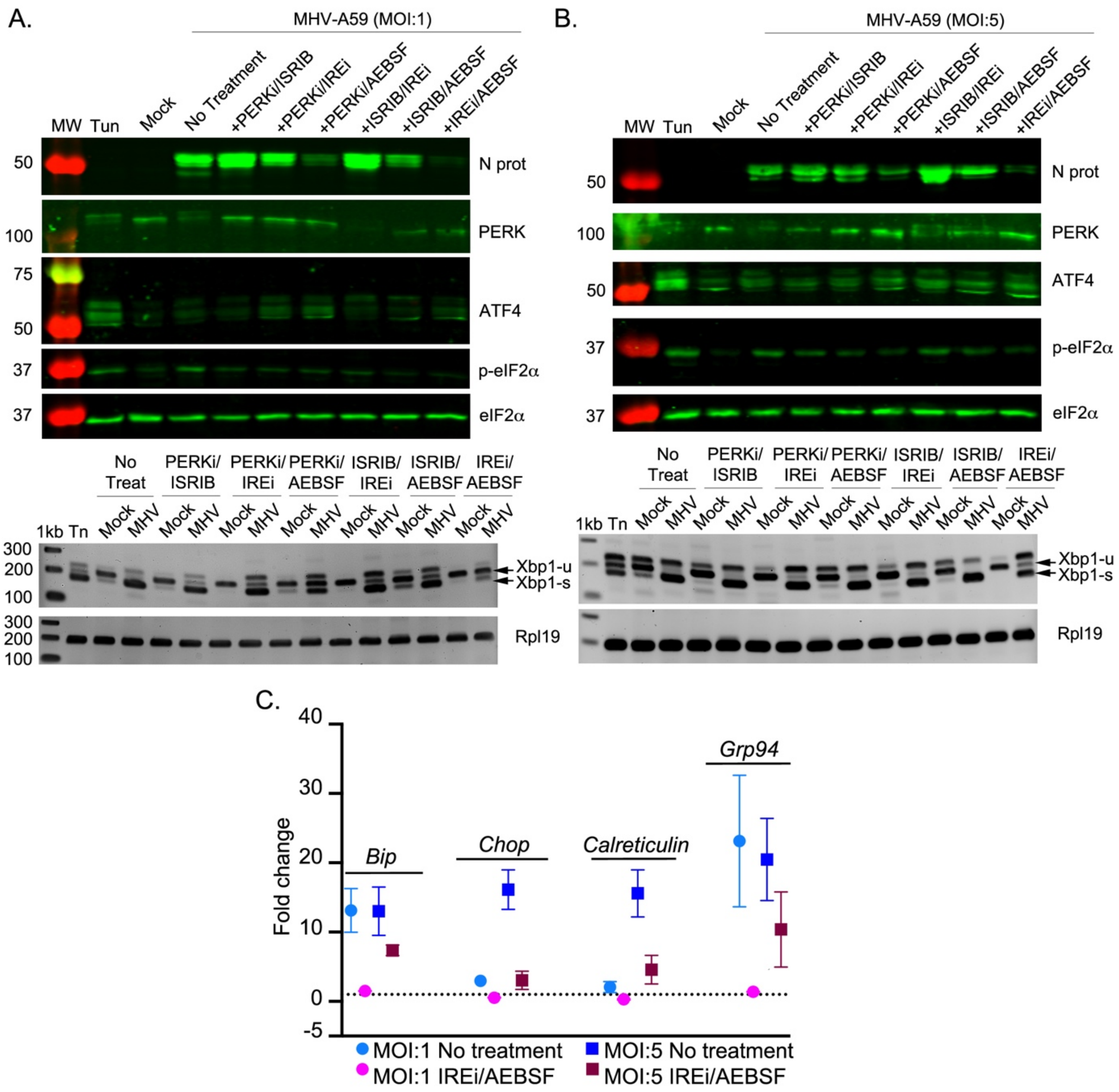


**S6 Figure. Effect of dual combinations of UPR inhibitors on activation of the three branches of the UPR during MHV infection.** MHV-infected cells were treated with the combinations of the UPR inhibitors shown in Fig 3B. The inhibitors were added to the cells immediately after the virus adsorption period and maintained in the medium until cells were harvested 8 h later. Cell lysates (upper) of MHV-infected cells at MOI 1 (**A**) and at MOI 5 (**B**) were immunoblotted using anti-N, anti-PERK, anti-ATF4, anti-p-eIF2α and anti-eIF2α as described in Fig 2A. The N protein panels have been duplicated from Fig 3C to facilitate comparison. RT-PCR analysis of *Xbp1-u* and *Xbp1-s* mRNAs (lower panels) as described in Fig 2D. (**C**) RT-qPCR of *Bip, Chop, Calreticulin* and *Grp94* mRNA for three biological replicates of a timecourse of MHV infection under no-drug or IREi 60μM/AEBSF 100μM treatment conditions as described in Fig 2B.

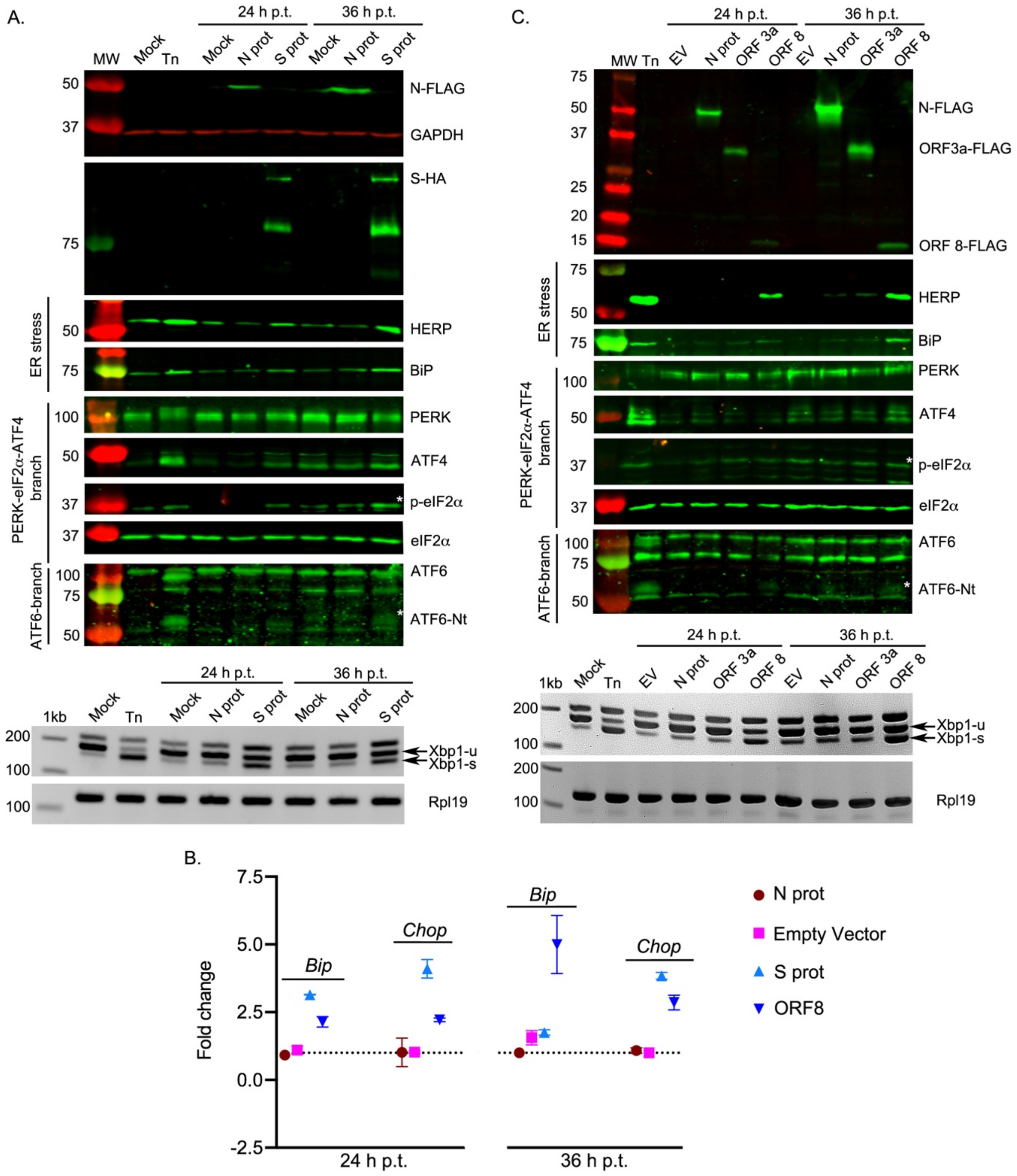


**S7 Figure. Mechanistic analysis of UPR activation by SARS-CoV-2 proteins.** (**A**) HEK-293T cells were transfected with plasmids encoding SARS-CoV-2 S (S-HA) or N (N-FLAG), mock-transfected, or treated with tunicamycin (Tn). Cells were harvested at 24 and 36 h p.t. and cell lysates (upper) were separated by 12% SDS-PAGE and immunoblotted using anti-FLAG, anti-HA, anti-HERP, anti-BiP, anti-PERK, anti-ATF4, anti-p-eIF2α, anti-eIF2α and anti-ATF6 as described in Fig 2A. The specific ATF6-Nt band is indicated with a white asterisk. RT-PCR analysis of *XBP1-u* and *XBP1-s* mRNAs (lower) as described in Fig 2D. (**B**) RT-qPCR of *BIP* and *CHOP* mRNA from two biological replicates of HEK-293T cells transfected with SARS-CoV-2 N, S and ORF8 or pcDNA.3 as an empty vector harvested at 24 and 36 h p.t. (**C**) HEK-293T cells were transfected with plasmids encoding SARS-CoV-2 ORF3a (ORF3a-FLAG), ORF8 (ORF8-FLAG), N (N-FLAG) or pcDNA.3 as an empty vector (EV) control, or mock-transfected, or treated with tunicamycin (Tn). Cells were harvested at 24 and 36 h p.t. and cell lysates (upper) and total RNA (lower) were analysed as previously indicated. "h p.t." = hours post-transfection.

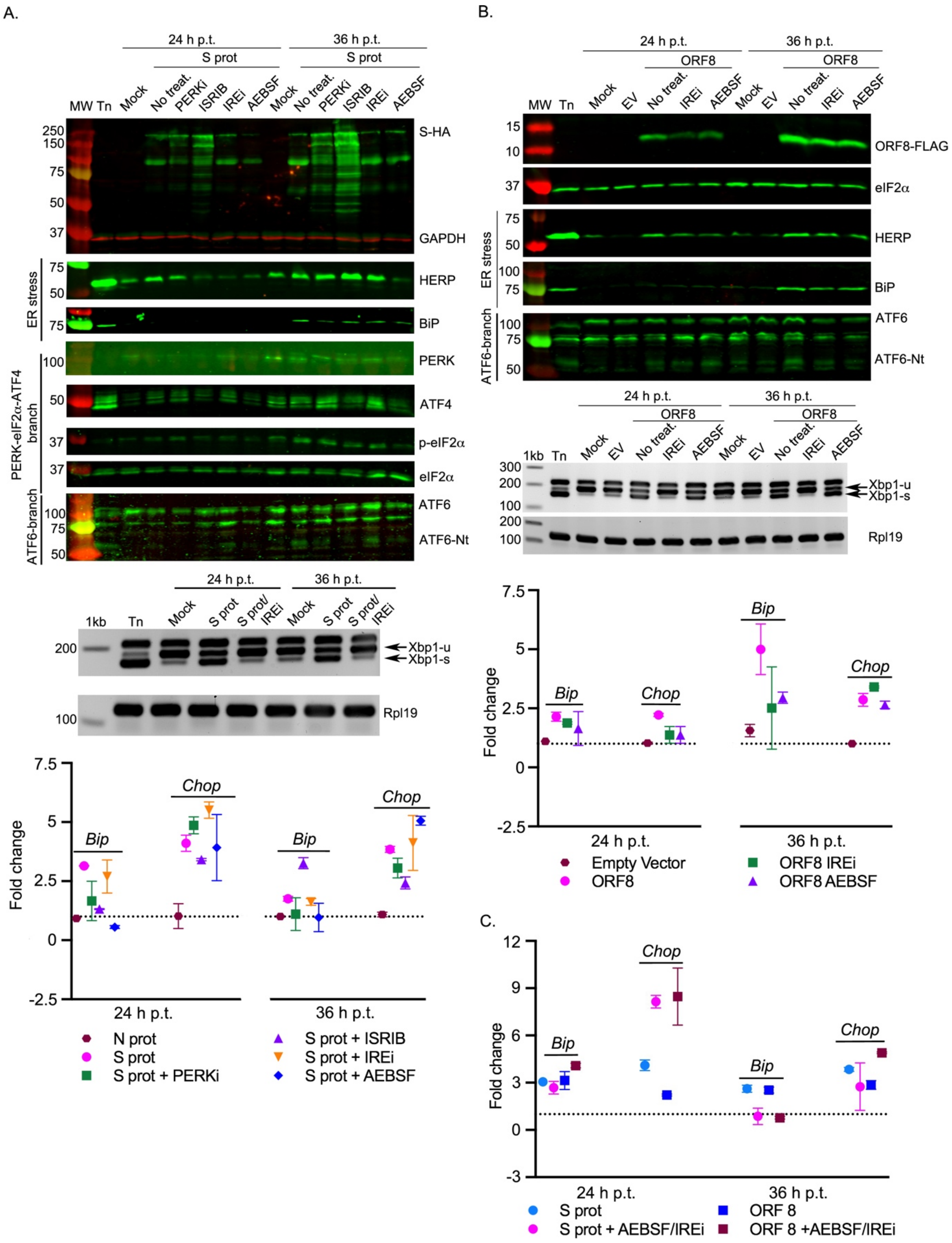


**S8 Figure. UPRi treatment reverses the activation of the UPR by SARS-CoV-2 proteins. (A)** HEK-293T cells were transfected with a plasmid encoding SARS-CoV-2 S (S-HA), mock-transfected, or treated with tunicamycin (Tn). At 8 h p.t., cells were treated with UPR inhibitors (5 μM PERKi, 2 μM

ISRIB, 60 μM IREi or 100 μM AEBSF) and then harvested at 24 and 36 h p.t. Cell lysates (upper) were immunoblotted using anti-HA, anti-HERP, anti-BiP, anti-PERK, anti-ATF4, anti-p-eIF2α, anti-eIF2α, anti-ATF6 and anti-GAPDH as described in Fig 2A. RT-PCR analysis using primers flanking the *XBP1* splice site (middle panel) as described in Fig 2D. RT-qPCR (lower) of *BIP* and *CHOP* mRNA from two biological replicates of HEK-293T cells transfected and treated as described in panel A. Data are normalised as described in Fig 2B. The specific p-eIF2α and ATF6-Nt bands are indicated with a white asterisk. (**B**) HEK-393T cells were transfected with a plasmid encoding SARS-CoV-2 ORF8 (ORF8-FLAG). At 8 h p.t., cells were treated with 60 μM IREi or 100 μM AEBSF and then harvested at 24 and 36 h p.t. Western blotting (upper), RT-PCR (middle) and RT-qPCR (lower) were performed as described in panel A. (**C**) RT-qPCR of *BIP* and *CHOP* mRNA from two biological replicates of HEK-293T cells transfected with SARS-CoV-2 S and ORF8 harvested at 24 and 36 h p.t. Cells were treated with 60 μM IREi and 100 μM AEBSF or a no-drug treatment control. Data are normalised as described in Fig 2B. RT-qPCR data of SARS-CoV-2 N, S, ORF8 and empty vector in panels A and B are reproduced from Supplementary Fig 7B for comparison to the treated conditions, which were performed as part of the same experiment. "h p.t." = hours post-transfection.

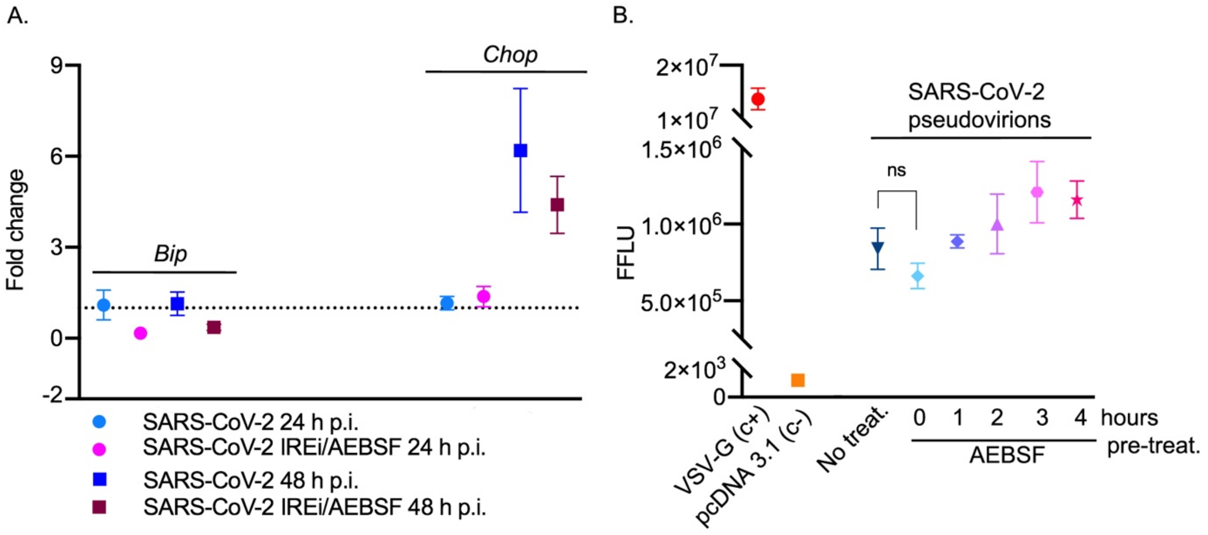


**S9 Figure. Effect of the UPR inhibitors on SARS-CoV-2 infection.** (**A**) RT-qPCR of *BIP* and *CHOP* mRNA from three biological replicates of Vero CCL81 cells infected and treated as described in Fig 5A. Data are normalised as described in Fig 2B. (**B**) Infectivity of lentiviral particles pseudotyped with SARS-CoV-2 S protein. HEK-293T cells were transfected with ACE2 and TMPRSS2 and treated with 100µM AEBSF between 0 and 4 h. Lentiviral particles engineered to contain a firefly luciferase reporter were pseudotyped with SARS-CoV-2 S, or vesicular stomatitis virus glycoprotein (VSV-G) as a positive control (c+, red) and empty pcDNA 3.1 vector as a negative control (c-, orange). Infectivity was measured as firefly luciferase units. Values show the mean averages of three biological replicates. Error bars represent standard errors. All *t*-tests are two-tailed and do not assume equal variance for the two populations being compared.

### Supplementary Tables

**S1 Table: RiboSeq and RNASeq library composition.** Number of reads assigned to each category. Reads under 25 nt long were designated "too short".

**S2 Table: Differential gene expression results**

**Sheets 1-4:** Ranked lists of genes which passed the thresholds of $\log_2$(fold change) greater than or equal to 1 (corresponding to a fold change of 2) and *p* value less than or equal to 0.05 after false discovery rate (FDR) adjustment for multiple testing. Sheets are as follows: TS_up, TS_down – genes which are significantly more (up) or less (down) transcribed in infected samples compared to mock, TE_up, TE_down – genes which are significantly more (up) or less (down) efficiently translated in infected samples compared to mock. Read counts for each sample, normalised by the total number of host-mRNA-mapping reads for that sample, are given in the right-most columns. **Sheets 5 and 6:** Full lists of genes which pass the threshold for inclusion in the analyses (requires ten reads mapping to this gene between all samples). TS = transcription, TE = translation efficiency.

**S3 Table: Reactome pathway and GO term enrichment analysis results**

**Sheets 1-4:** Enriched Reactome pathways. Lists of mouse gene names of significantly differentially expressed genes (Supplementary Table 2) were used for Reactome pathway enrichment[9], in which they were converted to their human orthologues and analysed to determine which pathways are significantly over-represented. Input gene lists are indicated in the sheet name, for example 'Reactome_TS_up' shows the Reactome enrichment results generated using the 'TS_up' list from Supplementary Table 2 as input. **Sheets 5-8:** Enriched GO terms. The same differentially expressed mouse gene lists were used for GO term enrichment analysis by PANTHER[101], against a background list of all the genes which passed the threshold for inclusion in that expression analysis. Column labels are as described in both Reactome and PANTHER user guides. All results with significant *p* values ($\leq 0.05$) are shown.

**S4 Table: List of genes classified as translationally resistant to eIF2α phosphorylation, based on Andreev *et al* [16]**. The first column shows the human genes classified as p-eIF2α-resistant by Andreev *et al* (excluding those from IRESite, which were not found to be p-eIF2α-resistant in their study). The list of genes used for the enrichment analysis, displayed in the second column, was generated by identifying mouse homologues of the human genes using the NCBI Homologene database [Database resources of the National Center for Biotechnology Information. *Nucleic Acids Research* **44**, (2016)]. Genes which were significantly more efficiently translated during MHV infection compared to mock infection are highlighted in bold.

### Supplementary Materials and Methods:

#### Immunofluorescence

Cells seeded on top of glass coverslips were fixed using ultra-pure, methanol-free formaldehyde (Polysciences) at a final concentration of 4% and permeabilised with Triton (0.3% diluted in PBS). Subsequently, cells were blocked with 5% FCS for 10 min prior to primary antibody incubation. Coverslips were probed with primary antibodies diluted in 1% FCS at the recommended concentrations for 1 h at room temperature, followed by incubation with secondary antibodies for 1 h in the dark. Secondary antibodies were donkey anti-rabbit or goat anti-mouse antibodies conjugated to Alexa Fluor 488 or Alexa Fluor 594 (ThermoFisher Scientific). Nuclei were counterstained with DAPI (1 µg/mL, Sigma-Aldrich) for 5 min. Coverslips were mounted onto microscope slides with Prolong Gold Antifade mountant (ThermoFisher Scientific) and allowed to cure overnight at room temperature. Cells were then imaged using an Evos FL Auto 2 microscope (ThermoFisher Scientific) with 60X oil-immersion objective. All images were exported as TIFF files.

**Metabolic labelling:** 17 Cl-1 cell monolayers were infected with MHV-A59 at a MOI of 5 PFU/cell. At 5 h p.i., cells were washed twice with PBS and labelled for 1 h in methionine-free DMEM supplemented with 125 µCi/ml [$^{35}$S] methionine. After this period, cells were harvested, washed twice with PBS and resuspended in lysis buffer (50 mM Tris pH 7.5, 100 mM NaCl, 5 mM EDTA, 0.5% NP40). Cell lysate aliquots were mixed with Laemmli's sample buffer to a final concentration of 1X and subjected to 10% SDS-PAGE followed by autoradiography.
