## Supplementary material for "Manipulation of the unfolded protein response: a pharmacological strategy against coronavirus infection": Supp Table 5

**Supplementary Table 5. List of oligonucleotides used.** Note *fwd* indicates forward primer, and *rev* indicates reverse primer.

| **Name** | **Sequence (5′-3′)** |
| --- | --- |
| *Rpl19* *fwd* (mouse) | ATGCCAACTCCCGTCAGCAG |
| *Rpl19* *rev* (mouse) | TCATCCTTCTCATCCAGGTCACC |
| *Bip* *fwd* (mouse) | CCTGCGTCGGTGTGTTCAAG |
| *Bip* *rev* (mouse) | AAGGGTCATTCCAAGTGCG |
| *Chop* *fwd* (mouse) | ACGGAAACAGAGTGGTCAGTGC |
| *Chop* *rev* (mouse) | CAGGAGGTGATGCCCACTGTTC |
| *Xbp1* *fwd* (mouse) | GAACCAGGAGTTAAGAACACG |
| *Xbp1* *rev* (mouse) | AGGCAACAGTGTCAGAGTCC |
| *Gadd34* *fwd* (mouse) | GACCCCTCCAACTCTCCTTC |
| *Gadd34* *rev* (mouse) | TCTCAGGTCCTCCTTCCTCA |
| *Calreticulin* *fwd (mouse)* | TGTTACCAAGGCTGCAGAGA |
| *Calreticulin* *rev (mouse)* | GGCCTCTACAGCTCATCCTT |
| *Grp94* *fwd (mouse)* | AGTCGGGAAGCAACAGAGAA |
| *Grp94* *rev (mouse)* | TCTCCATGTTGCCAGACCAT |
| *Human RPL19 fwd* | ATGTATCACAGCCTGTACCTG |
| *Human RPL19 rev* | TTCTTGGTCTCTTCCTCCTTG |
| *Human BIP fwd* | CGGGCAAAGATGTCAGGAAAG |
| *Human BIP rev* | TTCTGGACGGGCTTCATAGTAGAC |
| *Human CHOP fwd* | ACCAAGGGAGAACCAGGAAACG |
| *Human CHOP rev* | TCACCATTCGGTCAATCAGAGC |
| *Human XBP1 fwd* | TTACGAGAGAAAACTCATGGC |
| *Human XBP1 rev* | GGGTCCAAGTTGTCCAGAATGC |
